# Enhancing Photosynthesis under Salt Stress via Directed Evolution in Cyanobacteria

**DOI:** 10.1101/2024.08.27.610004

**Authors:** Zhenxiong Jiang, Khondokar Nowshin Islam, Malory Wolfe, Michael O’Connell, Dykia Williams, Ashley Florance, David J. Vinyard, Xiaohui Zhang, Maxwell Brenner, Andor J. Kiss, Xianhua Liu, Xin Wang

## Abstract

A key aspect of enhancing photosynthesis is improving its recycling kinetics, enabling swift resumption of photochemical quenching following environmental disruptions or stress. Salt stress exacerbates high light stress in cyanobacteria and leads to severe yield losses in crop plants. Genetic traits that confer salt tolerance without compromising photosynthetic performance are essential for improving photosynthesis under these conditions. Here we applied accelerated evolution in *Synechococcus elongatus* PCC 7942 by conditionally suppressing its methyl-directed mismatch repair system to obtain beneficial genetic traits for enhanced photosynthesis under salt stress. Screening over 10,000 mutants, we isolated eight strains with increased biomass or sucrose productivity under salt stress. Genome sequencing revealed an average of 8-20 single nucleotide polymorphisms (SNPs) or indels per genome. Notably, mutations in the photosystem II (PSII) reaction center D1 gene, resulting in the amino acid changes L353F, I358N, and H359N at the carboxyl terminus of the pre-D1 (pD1) protein, improve photosynthesis under salt and combined salt and light stress by potentially accelerating D1 maturation during PSII repair. Phylogenetic analysis of pD1 across cyanobacteria and red algae highlights the broad significance of these adaptive genetic traits, underscoring the importance of leveraging evolutionary insights to improve photosynthesis under stress or fluctuating environments.

## Introduction

Photosynthesis converts solar energy into chemical energy stored in biomass and is fundamental to life on Earth. The theoretical maximum of photosynthetic conversion efficiency from solar energy to biomass ranges from about 4-6% in plants to 9% in microalgae^1^. However, the average observed efficiencies are typically less than 1% in plants and up to 3% in algae cultivated in photobioreactors^2,3^, motivating substantial efforts to enhance photosynthetic conversion efficiency to increase crop productivity and develop carbon capture technologies^4-6^. Despite its importance, enhancing photosynthetic efficiency remains a challenging task, often complicated by evolutionary trade-offs developed over billions of years.

Key to enhancing carbon fixation and overall photosynthesis in natural settings is the improvement of photosynthesis recycling kinetics. Oxygenic phototrophs, which use tandem photosystems to drive the Calvin-Benson-Bassham (CBB) cycle, are nearly at their theoretical limit for photosynthetic energy conversion efficiency^1^. Environmental fluctuations frequently disrupt the balance between energy production and consumption^7^, leading to suboptimal photosynthetic energy conversion. Strategies such as accelerating the recovery of non-photochemical quenching (NPQ) have been shown to improve carbon fixation over a growing season thus enhancing crop yield^8^. Under abiotic stresses, plants also actively repress growth to maximize survival through stress-triggered cell signaling^9^. This stress-growth trade-off reduces photosynthetic efficiency^10,11^, leading to lower crop productivity. However, increasing evidence suggests that improved growth under stress is possible^12-14^, and such phenotypes can be explained by Pareto optimality^15^, where certain genetic traits confer the best tradeoff among multiple conflicting tasks.

Cyanobacteria possess unique genetic traits, such as the gene encoding a bifunctional fructose-1,6-bisphosphatase/sedoheptulose-1,7-bisphosphatase, which has been successfully engineered into plants to enhance photosynthesis and growth^16,17^. Recent directed evolution studies further identified new genetic traits for improved cyanobacterial growth under high light or combined high light and temperature stress^18,19^, expanding the potential to optimize photosynthesis by leveraging these beneficial traits. Traditional directed evolution methods, however, typically rely on a binary growth/no growth phenotype for isolating mutants, which limits their application to extreme or lethal stress factors. Additionally, long-term evolution experiments tend to optimize traits for stress response, often at the expense of reduced growth under normal conditions^18^. Salt stress inhibits photosystem II (PSII) repairs in cyanobacteria and is a common stress factor for plant growth^20,21^. Moderate salinity can lead to severe yield losses across a range of crop species^22^. Identifying beneficial genetic traits in response to sublethal salt stress necessitates methodologies capable of capturing the intricate dynamics of cellular responses to such stressors.

In this study, we employ *in vivo* accelerated evolution coupled with a high-throughput screening system to identify genetic traits that enhance photosynthesis under salt stress in cyanobacteria. We hypothesize that a vast array of beneficial genetic traits for photosynthesis improvement are often obscured by simple mutations such as single nucleotide polymorphisms (SNPs) in cyanobacterial genomes, which can be revealed through short-term directed evolution. In this proof-of-concept study, we have identified novel SNPs in the PSII reaction center D1 gene of *Synechococcus elongatus* PCC 7942 (hereafter *S. elongatus*), which are Pareto-front traits that simultaneously optimize photosynthesis and enhance resistance to both salt and combined salt and light stress. Phylogenetic analysis reveals that one of these mutations is commonly found across cyanobacteria and red algae, while others are rare or even absent in the database, suggesting broad significance of these identified mutations in adapting photosynthesis to environmental stresses. This methodology provides a powerful tool for uncovering novel genetic traits that enhance photosynthesis under stress or other environmental perturbations, with the potential to improve crop yields through translational research.

## Results

### Enabling mutagenesis by controlling nitrogen sources

The overall experimental design features directed evolution using a hypermutator strain, followed by high-throughput screening in 96-well microplates to identify mutants with enhanced carbon fixation under salt stress. Experimental evolution with *S. elongatus* can be time-consuming due to its relatively slow doubling time, which approximates 10 hours under optimal growth conditions^23^. Disrupting the *mutS* gene of the methyl-directed mismatch repair system (MMR) has been shown effective in inducing higher mutation rates in *S. elongatus* genome^19^. To enable accelerated evolution and facilitate downstream high-throughput screening, we sought to construct a hypermutator strain that can easily switch between hypermutation and normal states to effectively induce mutagenesis and preserve mutations. To achieve this, we adopted a previously published method to construct a *S. elongatus* hypermutator strain mSe0 (*mutS*0::*nirA*p), by placing the *mutS* gene under an inducible promoter P*_nirA_* controlled by nitrogen sources^24^. In doing so, the hypermutation state is induced in cells grown in BG11 medium supplemented solely with ammonium, and mutations are maintained by switching to BG11 medium exclusively supplemented with nitrate (**Fig. 1a**).

**Figure 1.**
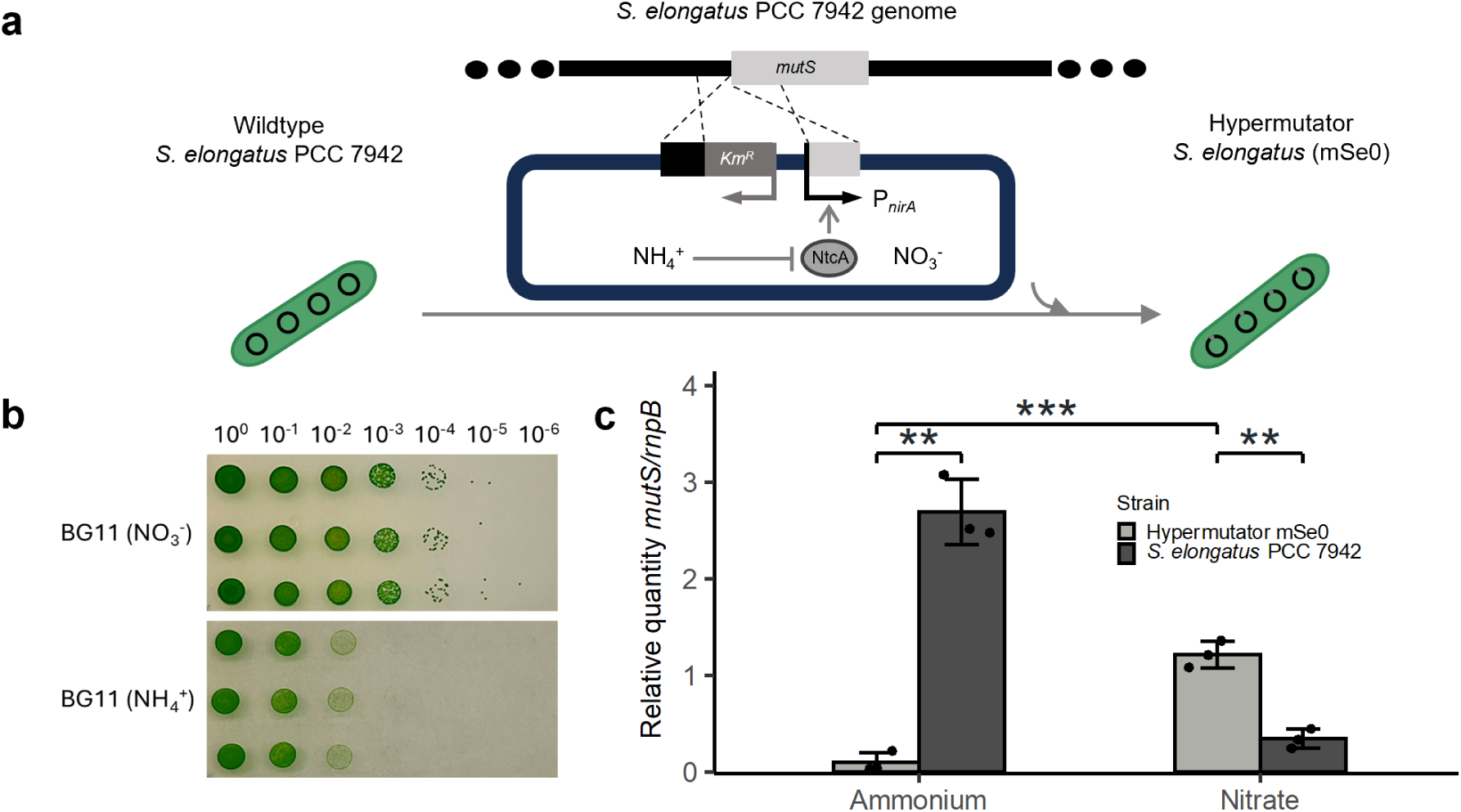
Construction of a hypermutator strain for directed evolution in *S. elongatus*. **a,** Schematic representation of mutagenesis controlled by nitrogen sources. Ammonium represses the *nirA* promoter by inhibiting its activator, NtcA. **b,** Spot assay. An equal amount of mSe0 cells (OD_730nm_ = 1) were diluted in triplicate onto BG11 agar plates supplemented exclusively with either nitrate or ammonium. Cyanobacteria in ammonium medium exhibit higher mutation rates, as evidenced by a reduced number of viable cells. **c,** RT-qPCR of *mutS* expression. RT-qPCR was conducted on cells cultivated for 24 hours in both BG11 (NO_3_^-^) and BG11 (NH_4_^+^) media. The relative expression of *mutS* was quantified against the reference gene *rnpB*. Error bars represent the standard deviation from three biological replicates, each with two technical replicates. Statistical significance was determined using a two-tailed Student’s t-test. ** *p* < 0.01, *** *p* < 0.001.

To confirm the higher mutation rate of the mSe0 strain, we first conducted a spot assay by diluting an equal amount of mSe0 cells onto BG11 medium supplemented exclusively with either nitrate or ammonium. Higher mutation rates increase the probability of hitting lethal genes, resulting in smaller viable cell populations in ammonium medium compared to nitrate (**Fig. 1b**). RT-qPCR further confirmed that the *mutS* gene expression in the mSe0 strain was reduced in ammonium compared to nitrate, and was lower than that in wild-type *S. elongatus* under the ammonium condition (**Fig. 1c and Supplementary Table 1**).

### Mutagenesis and high-throughput screening of cyanobacterial mutants with increased biomass and/or sucrose productivity under salt stress

The hypermutator strain underwent three rounds of mutagenesis experiments under the light intensity of 45 μE·m^−2^·s^−1^ in BG11 medium supplemented with increasing concentrations of NaCl (150 mM, 200 mM, and 250 mM). Each mutagenesis session was limited to four days, or approximately 8-10 cell generations, to carefully control mutation accumulation. This approach aims to generate potential Pareto-front genetic traits that enhance photosynthesis under salt stress without compromising normal growth performance (**Fig. 2a**). After each round, we grew mutants in BG11 nitrate medium supplemented with NaCl and screened for enhanced carbon fixation.

**Fig. 2.**
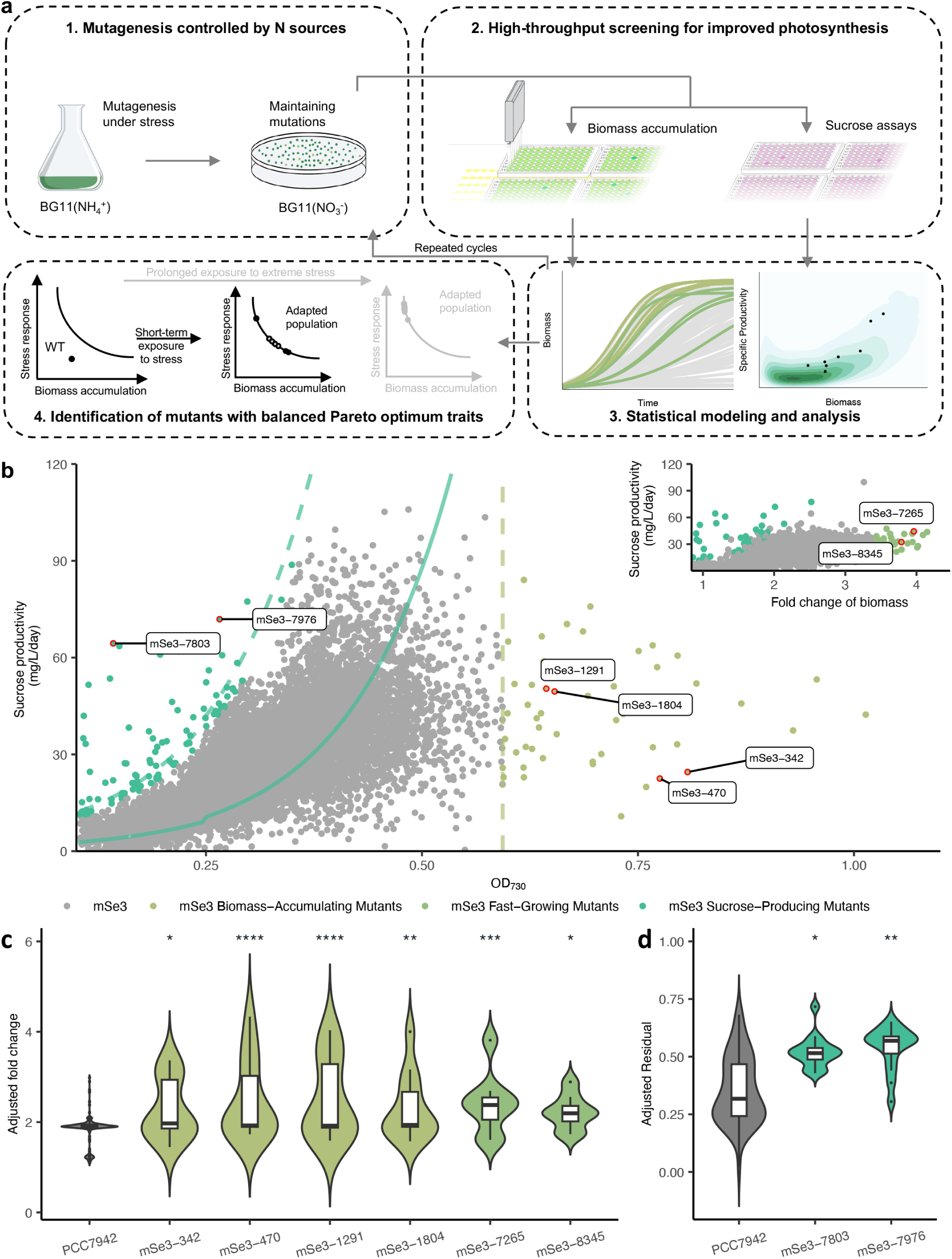
Short-term accelerated evolution to obtain Pareto-front traits for enhanced photosynthesis under salt stress. **a,** Overview of the high-throughput screening method. The hypermutator strain mSe0 is subject to three rounds of mutagenesis in BG11(NH_4_^+^) for a period of 4 days each round. Following each round, mutants are isolated on BG11(NO_3_^-^) agar plates to maintain mutations. Individual mutants are then subjected to high-throughput screening in 96-well microplates to assess both biomass (measured by optical density at 730 nm, OD_730nm_) and/or sucrose productivity. The final elite strain from each round serves as the starting point for the subsequent round of mutagenesis. The Pareto-front traits endow cyanobacterial mutants with diverse adaptive capabilities for biomass accumulation and stress response. **b,** High-throughput screening of mSe3 mutants. The top 1% of mutants with the highest standardized residuals of sucrose productivities were selected as the SPM candidates (green), and top 0.25% of mutants with the highest OD_730nm_ were selected as the BAM candidates (light green). A green solid line represents the fitted linear model based on the mSe0 control; The green dashed line represents two standard deviations (2σ) from the mean of sucrose productivity. The inset shows FGM candidates (medium green) representing the top 0.25% in population size fold change after a 54-hour incubation (24h normal growth followed by 30h growth with salt stress). **c**, Growth phenotype validation in 96-well cell culture plates. Population size fold change was normalized against the mean fold change of wild type *S. elongatus* for each batch to minimize the batch effects. The biomass validation with 12 replicates (4 replicates × 3 batches) reached a statistical power of 89.7%. **d**, Sucrose production validation based on 96-well cell culture plates. The validation with 16 replicates (8 replicates × 2 batches) reached a statistical power of 98%.

To effectively identify cyanobacterial mutants with enhanced photosynthesis under salt stress, we developed a high-throughput screening method to isolate desired mutants. The most reliable indicator of improved photosynthesis is an increase in terminal biomass, resulting from compounded higher carbon fixation over the cultivation period. Thus, we used biomass accumulation as one indicator of improved photosynthesis during the screening phase. In cyanobacteria, sucrose serves as a compatible solute for salt acclimation^25^. Sucrose is also the major photosynthesis end product in most plant leaves, with its synthesis closely linked to the rate of carbon fixation^26^. Therefore, we used sucrose accumulation as a second indicator for improved photosynthesis in *S. elongatus* mutants.

The first two rounds of mutagenesis, conducted under NaCl concentrations of 150 mM and 200 mM, served as preliminary screening steps to isolate potential candidate mutants. We utilized sucrose productivity as the sole quantitative metric for these rounds, selecting those mutants with the highest sucrose productivities within the population as final candidates. These candidate strains from each round were further engineered to secrete sucrose by incorporating the *Escherichia coli* sucrose permease (*cscB*) gene, as evidenced previously that increased sucrose production is closely linked to higher photosynthetic efficiency in the sucrose-secreting *S. elongatus*^27^. From the initial round of mutagenesis, approximately 700 mutants (mSe1) were screened, identifying mSe1-1086 as the final candidate strain (**Supplementary Fig. 1**). This strain served as the progenitor for the second round of mutagenesis under 200 mM NaCl, with approximately 1,200 mutants (mSe2) screened. Among these, the strain mSe2-842, exhibiting higher sucrose productivity compared to the sucrose-secreting *S. elongatus* control (**Supplementary Fig. 1**), was then selected as the starting strain for the subsequent round.

A larger-scale screening was conducted for the third round of mutants (mSe3) that underwent mutagenesis in BG11 supplemented with 250 mM NaCl. To increase the accuracy in isolating desired mutants, we optimized the inoculation and cultivation procedures to ensure that most mutants remained in the exponential growth phase during the high-throughput screening step (**Supplementary Fig. 2**). We further developed a regression model to quantitatively distinguish potential candidate mutants from controls based on sucrose productivity, using mSe0 as the baseline. This segmented linear model estimates the relationship between sucrose productivity and biomass for the control strain (**Supplementary Fig. 3**). Sucrose productivity of mSe3 mutants were then evaluated against this model to identify Sucrose-Producing Mutants (SPMs) with the highest 1% standardized residuals of sucrose productivities at different OD_730nm_. During the screening process, a subset of cells exhibited chlorosis. To assess its impact on sucrose productivity, we constructed a correlation analysis between different variables and chlorosis. The variables include terminal biomass (OD_730nm,54h_), terminal phycobilin level (OD_630nm,54h_), sucrose productivity, growth rate post-salt induction (OD_730nm,54h_/OD_730nm,24h_), and phycobilin pigment change rate (OD_630nm,54h_/OD_630nm,24h_). Correlation analysis showed no significant link between chlorosis and sucrose productivity (score of 0.28) (**Supplementary Fig. 4**), suggesting that chlorosis does not affect sucrose production. For biomass accumulation, the terminal biomass (OD_730nm,54h_) was used to determine potential Biomass-Accumulating Mutants (BAMs). Additionally, the biomass of a subset of mSe3 mutants was normalized to the initial inoculation density (OD_730,54h_/OD_730,0h_) to reflect their growth rates, with top candidates designated as Fast-Growing Mutants (FGMs). The inclusion of both terminal biomass and growth rate in our analysis is justified by the moderate correlation (score of 0.76) between the two (**Supplementary Fig. 4**), suggesting that each parameter captures distinct variables essential for identifying elite performers. In total, we screened over 10,000 mSe3 mutants, selecting the top 1% based on sucrose productivity as SPM candidates (**Fig. 2b**). Additionally, the top 0.25% based on terminal biomass (OD_730nm,54h_) were identified as BAM candidates (**Fig. 2b**), and another top 0.25% mutants based on biomass fold change (OD_730nm,54h_/OD_730nm,0h_) were selected as FGM candidates (**Fig. 2b** inset).

To validate the effectiveness of our models and identify elite mutants for either sucrose production or biomass accumulation, we conducted validation assays with sufficient replicates to achieve a statistical power of 90% (**Supplementary Fig. 5**). Using block ANOVA for comparison against the mSe0 control, we identified six mutants as the final elite BAM or FGM mutants with increased biomass accumulation, including mSe3-342, mSe3-470, mSe3-1291, mSe3-1804, mSe3-7265, mSe3-8345 (**Fig. 2c and Supplementary Fig. 5**). Additionally, two mutants, mSe3-7803 and mSe3-7976, were confirmed as elite Sucrose-Producing Mutants (SPMs) through block ANOVA (**Fig. 2d and Supplementary Fig. 5**). Importantly, these mutants exhibited similar photosynthetic rates to the *S. elongatus* wild type in batch cultures without stress (**Supplementary Fig. 6**), suggesting that potential mutations conferring salt tolerance did not compromise their normal growth performance. Interestingly, the mSe-342 strain, which was later discovered to exhibit a filamentous phenotype, failed to grow in the multi-cultivator, likely due to cell fragmentation caused by bubbling during growth.

### Mutations in BAM, FGM, and SPM elite strains

We sequenced all eight elite strains along with *S. elongatus* wild type with an average 1,000 × sequence coverage to identify potential beneficial mutations. As *S. elongatus* contains 3-4 genome copies during active growth^28^, we used a 0.25 allele frequency threshold to identify potential beneficial mutations, excluding the rest as potential sequencing errors or deleterious mutations.

Genome sequencing revealed mutations in *S. elongatus* wild type accumulated during domestication compared to the reference genome, including 19 missense/indel mutations and 10 synonymous mutations with a wide spectrum of allele frequencies (AFs). These loci were excluded from mSe3 strain mutations when identified with similar AFs. In total, 99 unique mutations were identified among all mSe3 strains with AFs > 0.25, with an average of 8-20 mutations found in each genome (**Fig. 3a**). Among these, 32 mutations are in intergenic regions and 67 mutations in coding sequences (CDS). The majority of the CDS mutations are either missense or indel mutations, with 10 being synonymous mutations (**Fig. 3a**, **Fig. 3b, and Supplementary Fig. 7**).

**Figure 3.**
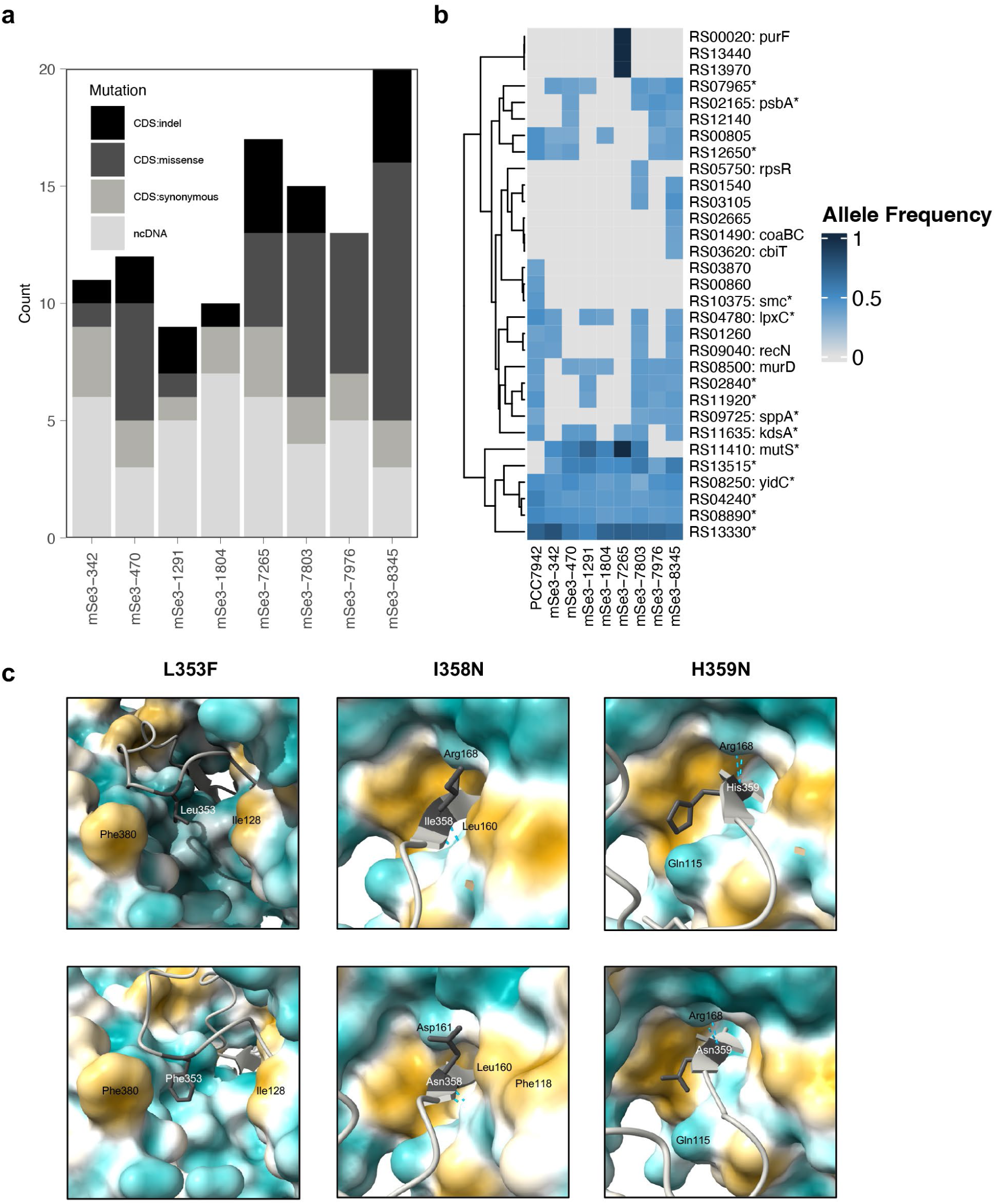
Mutation in BAM, FGM, and SPM elite mutants. **a,** Mutations from eight mSe3 strains and wild-type *S. elongatus* with allele frequencies above 0.25 are displayed. Mutations are classified into indels and SNPs in either coding sequences (CDS) or non-coding DNA (ncDNA) regions. **b,** Mutations located on CDS are shown, with shades of blue indicating allele frequency (AF). Grey shows sites with either the reference alleles or mutations with AFs of < 0.25. **c,** Interaction of pD1 and CtpA simulated by AlphaFold3^35^. CtpA is depicted using a surface representation, while the carboxyl terminus tail of wild-type pD1 (top row) and mutated pD1 (bottom row) are shown in ribbon format with mutated residues displayed in stick structures. Dark cyan represents hydrophilic surfaces, and dark goldenrod denotes lipophilic surfaces. Nearby amino acid residues around the mutation site are annotated.

Among the identified mutations, four elite mSe3 mutants (mSe3-470, mSe3-7803, mSe3-7976, and mSe3-8345) contain several mutations in the *psbA1* gene, encoding the core PSII D1 protein^29^ (**Fig. 3b and Supplementary Table 2**). The D1 protein is subjected to continuous photodamage and must be replaced every few minutes to ensure optimal photosynthesis in cyanobacteria and other oxygenic phototrophs^30^. Salt stress has been shown to inhibit transcription and translation of D1 protein during photodamage repair in *Synechocystis* sp. PCC 6803^21^. It is thus likely that mutations in the D1 gene might have led to improved growth under salt stress in the elite strains. The *psbA1* gene in *S. elongatus* encodes 360 amino acids, termed pre-D1 (pD1). During PSII assembly, the 16 amino acids at the carboxyl terminus are cleaved to form the mature D1 (mD1) by the protease CtpA^31^. Interestingly, all point mutations identified in the *psbA1* gene result in amino acid changes at the carboxyl terminus of the pD1 protein, leading to missense mutations at three positions, *i.e.* L353F, I358N, and H359N, either individually or in combinations (**Fig. 3c**). In cyanobacteria, pD1 processing is a two-step proteolytic process with the first cleavage step at the carboxyl terminus of Ala352 followed by the final cleavage at Ala344^32,33^. A previous study in *Synechocystis* sp. PCC 6803 further showed that Asn359 is crucial in preventing photoinhibition^34^. It is likely that these mutations near the proposed cleavage site or at the end of pD1 have led to more efficient cleavage during D1 maturation, resulting in longer duration of photochemical quenching and higher overall carbon fixation.

Among other identified mutations, we found an in-frame conversion of the start codon ATG to TTG in the *mutS* gene in several mutants, including mSe3-342, mSe3-470, mSe3-1291, mSe3-1804, and mSe3-7803, whereas mSe3-7265 had several mutations in the *mutS* coding region. These mutations might be results of survival strategies related to hypermutation. Many of the other identified CDS mutations are found in either hypothetical genes or obscure in their roles for photosynthesis improvement (**Fig. 3b**). We thus focused our efforts on validating the D1 gene mutations in potentially supporting improved photosynthesis in *S. elongatus*.

### Validating pD1 mutations toward tolerance to salt and light stress

High light results in the formation of reactive oxygen species and D1 damage, which is exacerbated by salt stress in cyanobacteria^21^. To test whether mutations in the *psbA1* gene confer beneficial traits for stress tolerance to salt and possibly light as well, we introduced an additional copy of the *psbA1* gene with either single mutations or their identified combinations into the neutral site III (NSIII) of the wild type *S. elongatus* genome, including *psbA1*^L353F^, *psbA1*^358N^, *psbA1*^H359N^, *psbA1*^T354P+I358N+H359N^, and *psbA1*^L353F+T354A+I358N^. Mutations at the position 354, specifically T354P/A were not found in individual mutants but were found in lower than 0.25 frequencies together with the other mutations. We thus included the T354P/A mutations in the validation experiment. A wild type *psbA1* gene copy was similarly introduced to the genome to serve as the control (*psbA1*^WT^). A mixture of native and heterologous pD1s will likely co-exist and be competitively recruited for PSII repair (**Fig. 4a and Supplementary Table 3**).

**Figure 4.**
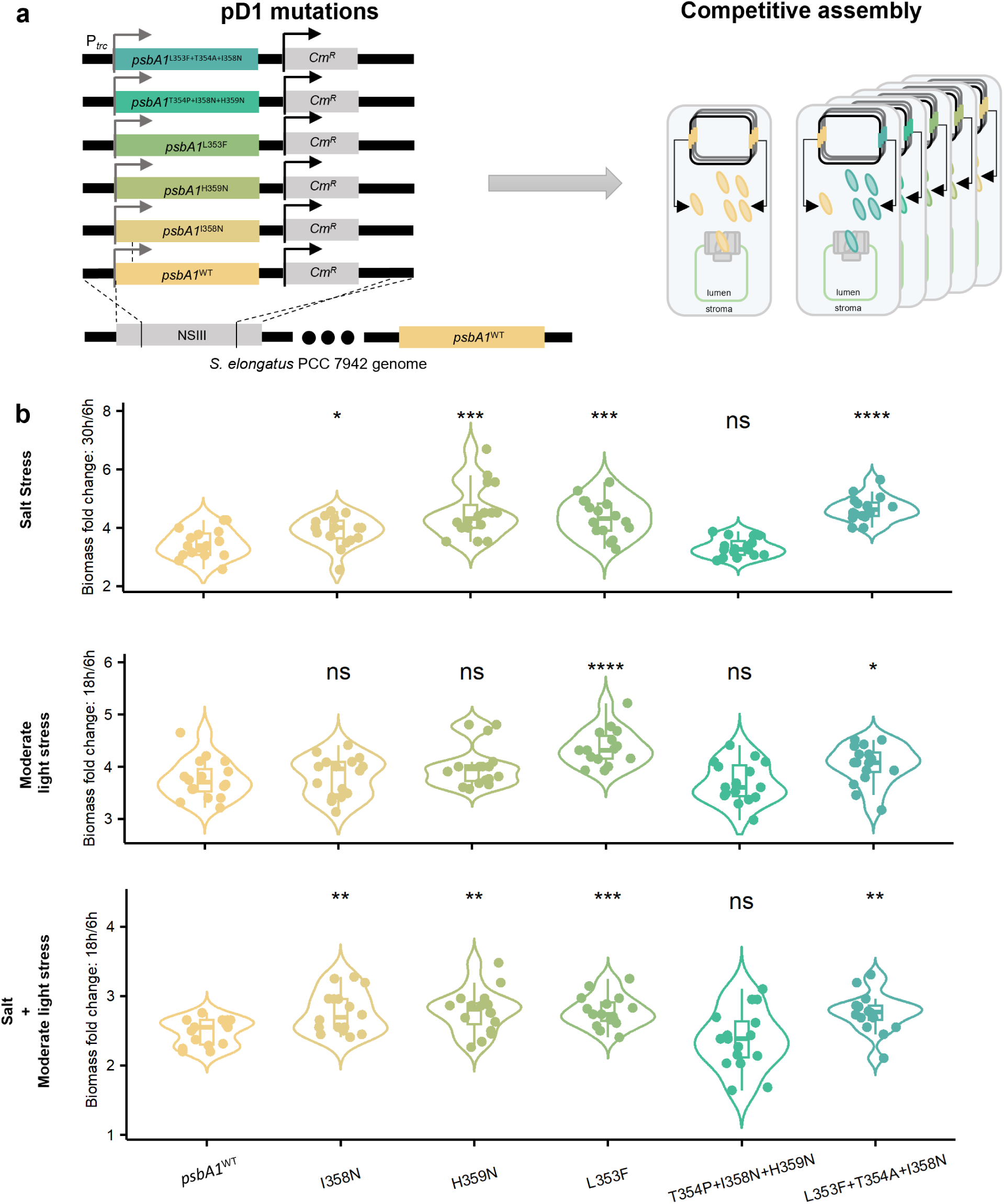
Validation of pD1 mutations under salt and moderate light stresses. **a,** Schematic representation of pD1 mutations and competitive assembly of D1 during PSII repair. **b,** Validation of stress resistance. Transformants with mutations L353F, I358N, H359N, and combined L353F+T354A+I358N showed superior biomass accumulation compared to the *psbA1*^WT^ control under 150 mM NaCl and combined stress of salt (150mM) and moderate light (110 μE·m^−2^·s^−1^) stress. Note that this light intensity leads to chlorosis of cyanobacteria in 96-well culture plates, suggesting moderate light stress. Transformants with L353F and combined L353F+T354A+I358N mutations outperformed the *psbA1*^WT^ control under 110 μE·m^−2^·s^−1^ light stress alone. Statistical analysis: Each experiment included 16 replicates per transformant, with statistical significance determined using an unpaired Student’s t-test (\**p* < 0.05, ** *p* < 0.01, *** *p* < 0.001, **** *p* < 0.0001).

We tested growth dynamics of these engineered D1 strains under salt stress (150mM NaCl) and elevated light conditions in both 96-well cell cultures (110 μE·m^−2^·s^−1^) and large batch cultures in multi-cultivators (300 μE·m^−2^·s^−1^). The pD1 mutations of L353F, I358N, H359N, and L353F+T354A+I358N all confer tolerance to salt stress and combined salt and light stress compared to the control (**Fig. 4b and Supplementary Fig. 8**). Interestingly, pD1 mutations of I358N and H359N do not seem to confer significant light tolerance on their own, suggesting salt tolerance as the dominant effect for their tolerance under combined salt and light stresses (**Fig. 4b and Supplementary Fig. 8**). In conclusion, L353F, I358N, and H359N in the pD1 protein confer tolerance to both salt and combined salt and light stresses in *S. elongatus*.

### L353F mutation of pD1 improves photosynthesis under elevated salt and light stress

To understand how photosynthesis is impacted by these pD1 mutations under salt and light stress in *S. elongatus*, we used the L353F mutation as an example to test for two considerations. Firstly, the mutation occurs immediately after the proposed first-stage cleavage site for pD1 carboxyl terminus^32^ (**Fig. 3c**), thereby could be involved in improved cleavage kinetics during D1 maturation. Secondly, the H359N mutation has been previously reported, showing its supporting role to prevent photoinhibition in *Synechocystis* sp. PCC 6803^34^. This suggests that asparagine near the end of pD1 could be important for PSII repair, as the I358N mutation also confers resistance to salt stress in our case (**Fig. 4**).

We grew both *psbA1*^WT^ and *psbA1*^L353F^ strains under the elevated light intensity of 300 μE·m^−2^·s^−1^ with three different NaCl levels (0 mM, 150 mM, and 300 mM) in multi-cultivators. The *psbA1*^L353F^ mutant exhibited similar growth to the *psbA1*^WT^ strain under 0 mM NaCl but showed increased growth under both 150 mM and 300 mM salt stress at the elevated light intensity (**Fig. 5a**). Full light spectrum scanning further revealed both increased chlorophyll (440 nm and 680 nm) and phycobilin (630 nm) contents in the *psbA1*^L353F^ mutant relative to the control (**Fig. 5b**), suggesting enhanced photosynthetic efficiencies under the combined salt and moderate light stress. The increase in chlorophyll was not as robust when light alone was used as the stress factor at three different light intensities (100, 300, and 1,500 μE·m^−2^·s^−1^) (**Supplementary Fig. 9**), indicating that L353F confers a trait beneficial for salt tolerance, with an added advantage under moderate light stress. Oxygen evolution rate measured at growing light intensities further confirms the increased photosynthetic efficiency of the *psbA1*^L353F^ mutant compared to the control under salt stress (**Fig. 5c**). When measured with saturating light intensity, the maximum photosynthetic efficiency of the *psbA1*^L353F^ mutant is also higher than the control under 150 mM NaCl, but this difference is diminished under the higher salt level of 300 mM (**Fig. 5d**). Taken together, these results demonstrate that L353F mutation endows both increased photosynthetic capacity and kinetics under salt and moderate light stress in *S. elongatus*, exemplifying a Pareto-front genetic trait for enhancing photosynthesis under sublethal stress conditions.

**Figure 5.**
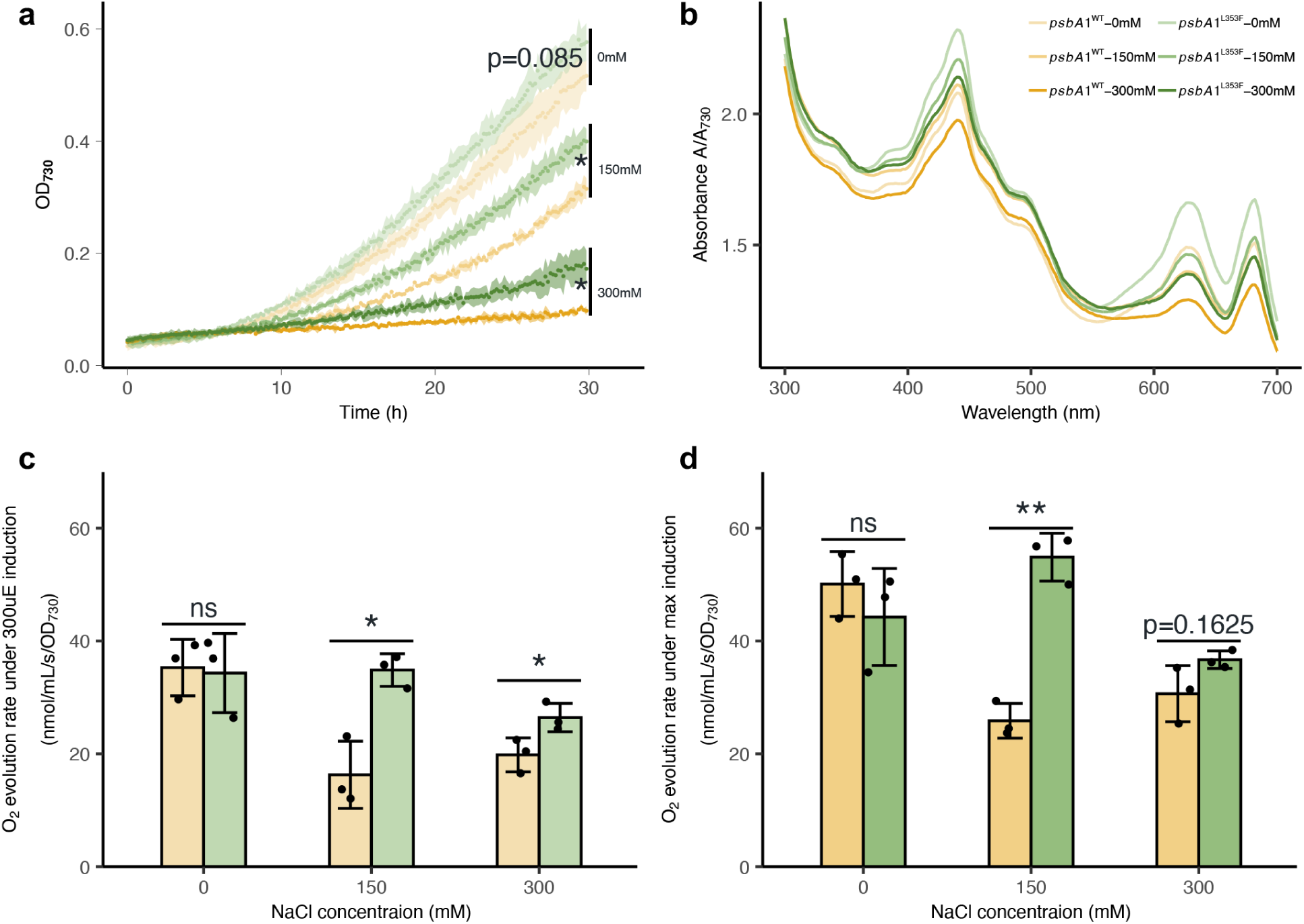
The pD1 L353F mutation confers improved growth and salt stress. **a,** Growth dynamics of *psbA1*^WT^ and *psbA1*^L353F^ transformants were monitored in triplicate in multi-cultivators. *psbA1*^WT^ is shown in yellow and *psbA1*^L353F^ in green, with increasing shading intensities corresponding to increasing salt concentrations (0mM, 150mM, 300mM). Data points represent the mean of three independent biological replicates, with ribbons indicating one standard deviation from the mean. Statistical analysis was performed using an unpaired Student’s t-test based on OD_730nm_ measurements at 30 h. **b,** Phycobilin content was estimated using full light spectrum scanning. Samples from late-log phase cells under different salt concentrations (0mM, 150mM, 300mM) and the light intensity of 300 μE·m^−2^·s^−1^ were analyzed in triplicate. Absorbance spectra ranging from 300 nm to 700 nm were recorded with a microplate reader, averaged from biological triplicates, and normalized to biomass (A_730_). Oxygen evolution rates were measured under growing light (**c,** 300 μE·m^−2^·s^−1^) and saturating light (**d,** 4,000 μE·m^−2^·s^−1^) conditions. Statistical significance was assessed using an unpaired Student’s t-test (\**p* < 0.05, ** *p* < 0.01).

### pD1 diversity in oxygenic phototrophs

Evolution has likely amassed a vast repertoire of beneficial genetic traits for adapting photosynthesis to various environmental conditions. We questioned whether the pD1 tail mutations observed through directed evolution in *S. elongatus* were present in the natural diversity of pD1 sequences. We assembled a database of 1,761 non-redundant D1 sequences from cyanobacteria, cyanophages, and red-lineage algae, all featuring a long pD1 carboxyl terminus tail of 16 amino acids. Only pD1 sequences that would result in PSII water oxidation activity were considered. The logo plot illustrates the observed natural variance in pD1 tail sequence (**Fig. 6**).

**Figure 6.**
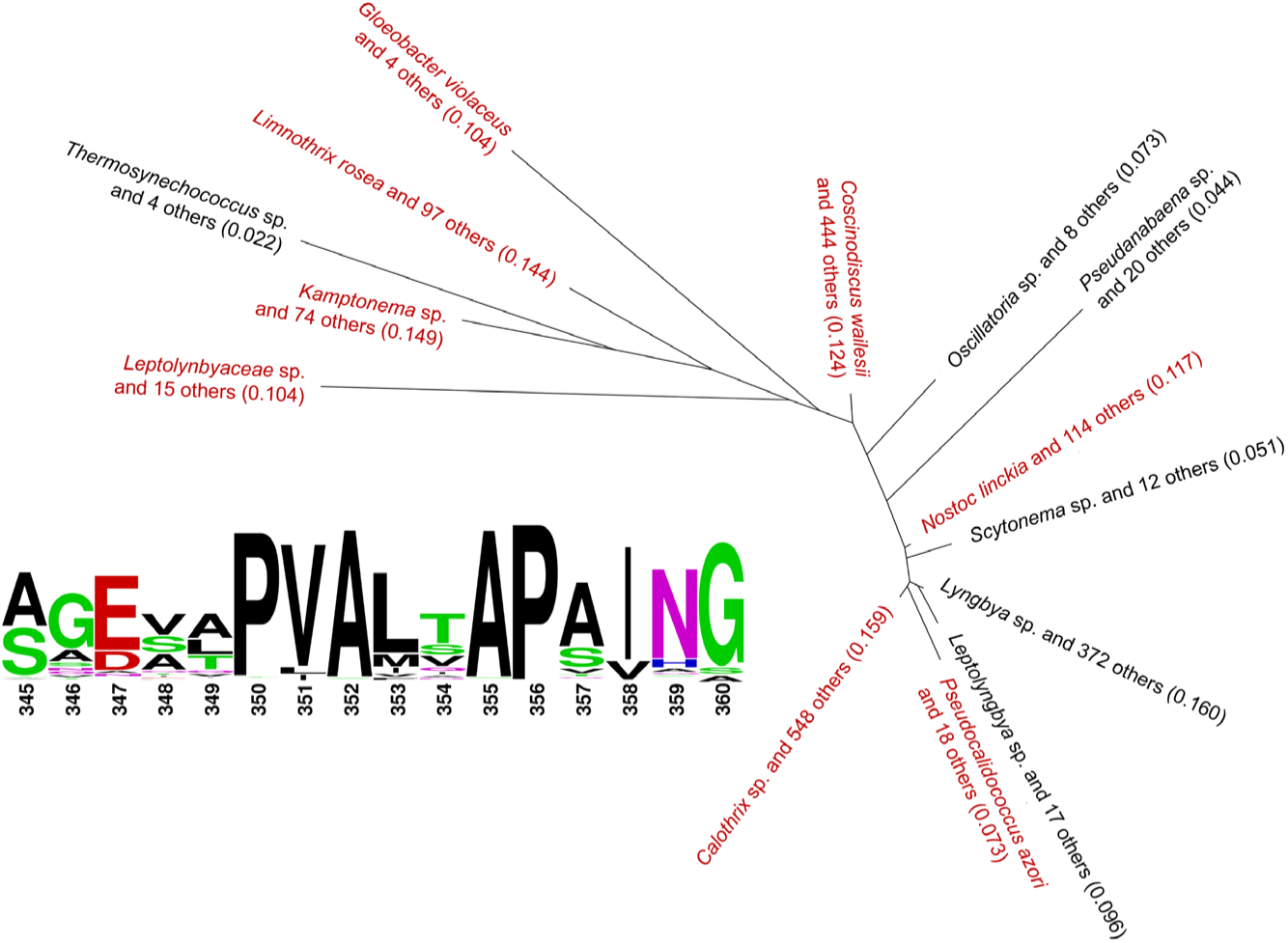
Phylogenetic analysis of pD1 sequences. Phylogenetic tree analysis and diversity of the pD1 tail were conducted using a database of 1,761 non-redundant, active D1 sequences from cyanobacteria, cyanophages, and red-lineage algae. A neighbor-joining phylogenetic tree based on the Jukes-Cantor model shows that F353 independently evolved multiple times among cyanobacteria and red algae. Subtrees with phylogenetic distance ≤ 0.15 were collapsed, with clades containing F353 in the pD1 tail sequences highlighted in red. The diversity within the 16 amino acid pD1 tail sequences is also depicted in a sequence logo, showcasing the evolutionary variability of this region.

This sequence analysis reveals a broad presence of the N359 across species, found in 70.8% of database sequences (**Fig. 6 and Supplementary Table 4**), highlighting the broad significance of the I359N mutation. Intriguingly, other mutations identified in this study are rare in nature. At position 353, hydrophobic residues such as L, M, and I are strongly favored. However, only 2.9% of data sequences contain F353, with phylogenetic analysis indicating its independent evolution across several clades (**Fig. 6**). Similarly, hydrophobic residues are preferred at position 358, making the I358N mutation notable as it is absent in all surveyed database sequences (**Supplementary Table 4**). These findings highlight the unique salt-tolerance feature of L353F and I358N pD1 mutations, underscoring the effectiveness of our method in identifying unique mutations that expand the evolutionary genetic repertoire for photosynthesis improvement under stress.

## Discussion

Cyanobacteria emerged over three billion years ago with the development of tandem photosystems to harvest solar energy, leading to the Great Oxidation Event on Earth^36^. With its current design, photosynthesis operates near its theoretical limit in converting solar energy into chemical energy stored in biomass^1^. An effective strategy to improve photosynthesis for emerging societal challenges, such as increasing crop yield, is to enhance its recycling kinetics to boost effective photochemical quenching under fluctuating environments^37^. Over billions of years, evolution has likely accumulated a vast repertoire of beneficial genetic traits among oxygenic phototrophs to adapt photosynthesis to constantly changing environments^38,39^, with potential Pareto-front genetic traits distributed throughout their genomes. Glimpses of these beneficial genetic traits can be seen embedded in the genomes of cyanobacteria via a few SNP mutations, as evidenced by endowing *S. elongatus* with a high-light-tolerant phenotype through complementing SNPs found in its close relative, *S. elongatus* UTEX 2973^40^. Recent directed evolution studies further validate this hypothesis, showing that new SNP traits can be revealed in model cyanobacteria for enhancing photosynthesis under high light or temperature stresses^18,19^. However, a limitation of applying directed evolution to enhance photosynthesis is its dependence on applying lethal stress factors, thus a binary growth/no growth phenotype can facilitate the isolation of potential beneficial mutants. To mitigate this challenge, we applied short-term *in vivo* accelerated evolution by conditionally suppressing the *mutS* gene of the MMR system, thereby easily inducing and maintaining SNP mutations for improved photosynthesis under salt stress (**Fig. 1**). Combined with a relatively high-throughput screening system, we were able to isolate eight *S. elongatus* mutants with enhanced biomass or sucrose productivity under salt stress by screening just over 10,000 mutants (**Fig. 2**), demonstrating the effectiveness of this approach in generating beneficial SNP mutations for improved photosynthesis.

The final elite strains contain an average of 8-20 SNPs and indels in their genomes, with the majority being missense mutations in CDSs and substitutions in non-coding regions (**Fig. 3b and Supplementary Fig. 7**). The mutations found in the *psbA1* gene, encoding the PSII reaction center D1 protein, were identified as the contributing factor for improved photosynthesis in the mutants. Intriguingly, these mutations were located at the carboxyl terminus of the pD1 protein at positions 353, 358, and 359. The 353 position is adjacent to the early cleavage site Ala352 during pD1 processing^32,33^, suggesting its potential involvement in altering pD1 cleavage efficiencies. In *Synechocystis* sp. PCC 6803, replacing Asn359 with either His or Asp resulted in higher vulnerability to photoinhibition^34^. In *S. elongatus*, there are three different copies of *psbA* genes in the genome. The *psbA1* gene encodes the D1 with His359 whereas *psbA2/3* genes encode the D1 with Asn359^41^, with *psbA2/3* gene expression induced under high light conditions. These previous findings suggest that Asn359 might be involved in photoprotection under high light or fluctuating light conditions. Interestingly, the mutation at position 359 in our study showed the conversion of His359 to Asn359 in the *psbA1* encoded D1, likely improving the D1 maturation process for robust photosynthesis under stress.

Mutations of L353F, I358N, and H359N in the pD1 protein confer stress tolerance to salt and combined salt and moderate light stress in *S. elongatus* (**Fig. 4**). Further validation using L353F as an example demonstrates both improved photosynthetic kinetics and maximum capacity due to the introduced mutation (**Fig. 5**). Previous studies in cyanobacteria and algae showed that mutants without the carboxyl terminus extension in D1 have normal PSII activity and growth but compromised competitiveness when co-cultured with the wild type^42-44^. A later study observed an increase in unassembled D1 protein and a slight increase in sensitivity to photoinhibition in the mutant strain lacking the C terminus extension, suggesting that this region could mediate D1 assembly into PSII^34^. Additionally, altering the C terminus extension could impact its binding efficiency with CtpA, thus affecting D1 maturation^31^. These previous studies indicate a strong association between pD1 carboxyl terminus diversity and robust photosynthesis. Follow-up studies on pD1 kinetics could help better clarify the role of the mutation in supporting photosynthesis.

Phylogenetic analysis of pD1 reveals a rare occurrence of F353 and even absence of N358 but a consensus for N359 at the carboxyl terminus in cyanobacteria, cyanophages, and red-lineage algae (**Fig. 6**), suggesting that asparagine at this position may offer adaptive benefits for photosynthesis under various environmental stresses. The D1 protein is highly conserved across kingdoms with consensus sequences in cyanobacteria and plants^45,46^. A single copy of the D1 gene is encoded in the chloroplast genome in plants. A recent study engineered an extra copy of the *psbA* gene in the nucleus genomes of *Arabidopsis*, tobacco, and rice, resulting in significantly enhanced carbon assimilation rates and increased biomass and grain yield^47^. Our identified beneficial mutations in cyanobacterial pD1 could thus lead to translational research by modifying plant pD1 for crop yield improvement. Our study showcases the advantage of applying short-term directed evolution to improve photosynthesis under salt and moderate light stress, providing a powerful tool to bring about novel beneficial genetic traits to adapt photosynthesis under various stress conditions.

## Materials and Methods

### Strains and culture condition

The wild-type *S. elongatus* PCC 7942 was cultured on BG11 agar plates for isolation and maintained in BG11 liquid media at 25°C under a light intensity of 40 μE·m^−2^·s^−1^ for routine cultivation. The *nirA* promoter sequence was amplified from *S. elongatus* wild type and used to construct pXWO3_nirA plasmid using NEBuilder HiFi DNA Assembly Master Mix. The mSe0 mutator strain, created from pXWO3_nirA transformation, along with mutants derived from directed evolution, were kept in BG11(NO_3_^-^) supplemented with 5 μg/mL kanamycin. During directed evolution experiments, the mutator strain was grown in BG11(NH_4_^+^) supplemented with 10 mM NaHCO_3_, 5 μg/mL kanamycin, and different concentrations of NaCl (150, 200, 250 mM). *S. elongatus* harboring the sucrose permease gene *cscB* was propagated on BG-11 agar plates containing 5 μg/mL chloramphenicol and 5 μg/mL kanamycin and cultivated in BG-11 liquid containing the same concentrations of antibiotics. *Escherichia coli* DH5α, which carries the NSIII *cscB* plasmid gifted from the Ducat lab, was cultured in LB broth with 25 μg/mL chloramphenicol and maintained on LB agar plates supplemented with 50 μg/mL chloramphenicol. **Plasmid transformation into *S. elongatus* strains.** *S. elongatus* cultures were grown to an OD_730nm_ of 1.0. One milliliter of each culture was collected and centrifuged at room temperature for 2 minutes at 13,000 rpm. The cell pellet was washed with 1 mL of 10 mM NaCl and centrifuged again. The resulting pellet was then resuspended in 200 µL of BG11 (NO_3_^-^) medium. To this suspension, 100 ng of plasmid DNA was added and gently mixed. The mixture was incubated in the dark at 30°C for 4-16 hours with gentle agitation. After incubation, the cultures were plated on BG11 (NO_3_^-^) agar plates containing the appropriate antibiotics and incubated at 30°C under light conditions.

### Reverse Transcription-Quantitative PCR (RT-qPCR)

*S. elongatus* wild type and mSe0 strains at mid-exponential growth phase were harvested by centrifugation. The cell pellets were washed and resuspended in either BG11 (NO_3_^-^) or BG11 (NH_4_^+^) medium. Biological triplicates of both strains were then inoculated in their respective media and cultured to mid-exponential phase at 30°C under continuous light at 45 μE·m^−2^·s^−1^. For RNA isolation, cultures were collected and centrifuged at 4°C, and RNA was extracted using the TRIzol-chloroform-isopropanol method according to the manufacturer’s protocol (Fisher Scientific, Pub. No. MAN0001272). RNA was subsequently reverse-transcribed to cDNA using the iScript cDNA Synthesis Kit (Bio-Rad, INST-653 Ver D). For the qPCR analysis, primers for *mutS* and the reference gene *rnpB* were designed using Primer3 (**Supplementary Table 1**). Standards and samples were run in technical duplicates in the CFX 96 Connect Real-Time PCR system (BioRad) following the SYBR Green kit protocol. The differential gene expression of *mutS* was measured by qPCR using CFX Connect software 3.1 employing the ^ΔΔ^Cq method relative to the ‘reference’ gene *rnpB*.

### Spot assay

mSe0 mutants at early-mid log phase were resuspended to an OD_730nm_ of 1 in BG11(NO_3_^-^) and BG11(NH_4_^+^), respectively. After 1:10 serial dilution, 5 µL aliquots were spotted onto corresponding agar plates supplemented with kanamycin. Plates were incubated at 30 °C under a light intensity of 40 μE·m^−2^·s^−1^ for one week. The survival colonies with the same dilution are results of different mutation rates between nitrate and ammonium conditions.

### Directed Evolution

A single colony from the mSe[i] strain series was grown in BG11(NO_3_^-^) in a 6-well plate at 30 °C and 30 μE·m^−2^·s^−1^ until visibly green. The culture was then scaled up in flasks (1:20 v/v) and grown until the OD_730nm_ reached 0.7-1.0. Cells were harvested by centrifugation at 10,000 rpm for 1 minute at room temperature, washed three times with BG11(NH_4_^+^), and resuspended in BG11(NH_4_^+^) for mutagenesis in a 6-well plate. Cultivation continued for 4 days in a shaking incubator set to 230 rpm, 30 μE·m^−2^·s^−1^ light intensity, and 30°C, with additions of kanamycin and NaCl. Following mutagenesis, cells were washed three times with BG11(NO_3_^-^), resuspended in 100 µL of BG11(NO_3_^-^), diluted appropriately, and plated on BG11(NO_3_^-^) agar plates supplemented with kanamycin.

### Optimization of mSe0 growth in 96-well microplates

The mSe0 strain was initially grown in 20 mL of BG11(NO_3_^-^) until its OD reached approximately 0.1. Subsequently, 1 µL of this culture was transferred into each well of a 96-well microplate containing 249 µL of BG11(NO_3_^-^) supplemented with 5 µg/mL kanamycin. The microplates were incubated at 30°C under continuous light at an intensity of 45 μE·m^−2^·s^−1^. Optical density at 730 nm was measured hourly to monitor growth. At 24 hours, 25 µL concentrated NaCl stock in BG11 (NO_3_^-^) was added into culture for salt induction, and hourly data collection continued until 54 hours. The growth rates before and after salt induction were analyzed using an exponential growth model. The optimized bacterial growth was depicted in Supplementary Figure 2.

### High-throughput assay of biomass and sucrose productivities

Mutant colonies were maintained on agar plates and transferred to 96-well plates, each well containing 250 µL of BG11(NO_3_^-^). Growth and salt induction were conducted following the optimized procedure above. Optical densities at inoculation (OD_730nm,0h_), salt induction (OD_730nm,24h_), and post incubation (OD_730nm,54h_) were measured using a Molecular Devices SpectraMax® iD5 Multi-Mode Microplate Reader. Post incubation, cells were subjected to sucrose assay using the Eppendorf epMotion® 5073m liquid handling workstation. Cyanobacterial cells are lysed with the addition of 25 µL of lysis solution (2% dodecyltrimethylammonium bromide (DTAB) in 0.4 M NaOH) per 50 µL sample, followed by neutralization with 0.4 M HCl after 5 minutes of shaking at 1,100 rpm. Post lysis, 50 µL of invertase solution was added, incubated at 50 °C for 1 hour, followed by the addition of 100 µL of Megazyme GOPOD reagent and a further 20 minute incubation at 50°C. OD_510nm_ was then measured, and sucrose concentration was calculated against the standard curve ranging from 0.05 mg/mL to 0.1 mg/mL. Sucrose productivity is calculated by incorporating growth rate before (*r_1_*) and after (*r_2_*) salt induction. Given the exponential growth model, the following formula was used to calculate sucrose productivity:

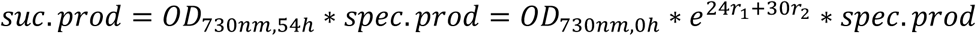

This method captures dynamic growth changes influenced by environmental conditions, facilitating a more accurate assessment of the mutants’ sucrose productivity relative to the growth.

### Identification and validation of candidate strains from high-throughput screening

To identify potential candidate strains with increased sucrose productivities, the mSe0 strain was used to construct a model to estimate the relationship between sucrose productivity and biomass. To do this, mSe0 was randomly inoculated into 96-well plates across a spectrum of OD_730_ values. Sucrose and growth assays were performed as described above. The obtained data formed the training set, which was first grouped using k-mean clustering, followed by analyzing their relationships using a linear regression-based model. This model used log-transformed sucrose productivity as the response variable, and terminal OD_730_ as the independent variable:

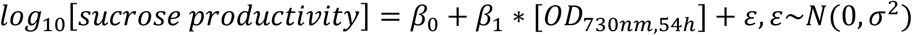

Sucrose-producing candidates were selected based on the highest 1% standardized residuals of sucrose productivities at different OD_730nm_. Additionally, mutants in the top 0.25 % of both terminal biomass (OD_730nm,54h_) and biomass fold change (OD_730nm,54h_ /OD_730nm,0h_) were selected as biomass-accumulating candidates and fast-growing candidates, respectively.

To identify elite mutants with statistical significance compared to the control strain mSe0, power analysis was used to establish an appropriate sample size, ensuring a power of 0.9 and an alpha threshold of 0.05. Assays for sucrose productivity, growth rate, and biomass accumulation were repeated following initial screening procedures. Statistical analyses were performed using block ANOVA, with ‘block’ representing different experimental batches.

### Correlation analysis between chlorosis and multiple variables

To assess the impact of chlorosis on sucrose production and growth experienced by some mutants during high-throughput screening, Pearson correlation analysis was performed. The chlorosis status of mutants was first determined using a classifier trained on a dataset including mSe0 and a subset of mSe3 cells. The variables include sucrose productivity, OD_730nm,54h_, growth rate 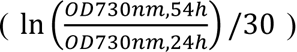, OD_630nm,54h_, and phycobilin change rate post salt induction 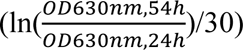. All predictors were standardized by their means and standard deviations to ensure comparability.

### Genome extraction and sequencing

The cyanobacterial culture was grown in BG11(NO_3_^-^) supplemented with 20 mM NaHCO_3_ and 5 µg/mL kanamycin. Cells were harvested for genome extraction when OD_730nm_ reached between 0.5 and 0.6. The Promega Wizard® HMW DNA Extraction Kit was employed for genomic DNA (gDNA) extraction, using approximately 3 × 10^9^ cells. The extracted genomic DNA, with yields exceeding 1.2 µg, was utilized for PCR-free library preparation and subsequent genome sequencing by Novogene with an average of 1,000 × genome coverage based on 150 bp paired-end reads.

### Genetic variant calling and mutation analysis

To ensure the sequencing quality, all FASTQ files were inspected using FastQC v0.11.9^48^. Low-quality bases in each FASTQ file were later trimmed using Trimmomatic v0.39^49^. Adapters and low-quality reads including leading and trailing bases below quality 3 or N bases were removed. A 4-base window was used to slide and scan all reads and cut reads if average quality was less than 15. At the end, remaining reads shorter than 36 bases were discarded. The trimmed reads were then mapped against the *S. elongatus* PCC 7942 reference genome from NCBI (Assembly: GCF_000012525.1) using Bowtie2 v2.2.5 with default parameters^50^. Variant calling for SNPs and indels was performed using samtools and bcftools (version 1.9)^51^. The process involved converting SAM files to BAM format, sorting the BAM files by genomic location, and converting sorted BAM files to text pileup output. Specifically, no anomalous read pairs were skipped, and both base and mapping quality thresholds were set to 0. In addition, several annotation information was added including allelic depth (AD), allelic depths on the forward strand (ADF), allelic depth on the reverse strand (ADR), number of high-quality bases (DP), Phred-scaled strand bias P-value (SP). bcftools call was then used to obtain mutations. All alternative alleles present in the alignments even if they did not appear in any genotypes were collected, multiallelic caller was adopted due to the polyploidy of cyanobacteria. Genotype quality (GQ) was assessed for calling quality. The resultant VCF file underwent annotation with snpEff v5.2 using default settings, and annotated VCF files were tabularized using the snippy-vcf_to_tab function in the snippy package (v4.6.0)^52^. A customized R script was used to perform all allele frequency analysis.

### Mutation complementation

The *psbA1* mutations were introduced in reverse primers (**Supplementary Table 4**) and amplified with a universal forward primer for constructing all complementing mutant strains. The amplified *psbA1* genes were purified and assembled using NEBuilder HiFi DNA Assembly Master Mix, and the constructs were transformed into *S. elongatus* PCC 7942 wild type. New colonies were initially grown on low-phosphate BG11 supplemented with 5 μg/mL chloramphenicol for two days to enable faster segregation, then transferred to standard BG11 plate supplemented with 5 μg/mL chloramphenicol. Segregation was verified by PCR amplification of the NSIII region. Transformants were cultured in 24-well plates for preculture, then transferred to 96-well plates containing BG11(NO_3_^-^) with 20mM NaHCO_3_ with or without 150mM NaCl. Sixteen replicates per mutant per condition were inoculated to ensure statistical robustness. Mutants were exposed to a light intensity of 110 μE ·m^−2^ ·s^−1^ (moderate light stress) and/or 150 mM NaCl to assess growth. OD_730nm_ readings were taken at 0h, 6h, and either 18h or 30h depending on whether salt was added. For batch cultures, mutants were grown in multi-cultivators at 100 μE ·m^−2^ ·s^−1^ and 30 °C, harvested at mid-log phase, and inoculated in triplicate into new media at a starting OD_730nm_ of 0.05. Growth was monitored under different light conditions (100, 300, 1,500 μE ·m^−2^ ·s^−1^) and salt concentrations (0, 150, 300 mM NaCl).

### Oxygen evolution and full light spectrum absorbance measurements

During mid-log phase, cell cultures were sampled for oxygen evolution using a Hansatech Oxytherm+ system. After a 10-minute dark incubation, oxygen evolution was measured for 7 minutes either at growing light intensities or saturating light intensity using the built-in LED panel, followed by the measurement of respiration immediately after termination of illumination. Oxygen evolution rates were normalized to biomass OD_730nm_. For absorbance measurements, biological triplicates with three technical replicates of 250 µL cell culture were analyzed in a BioTek Synergy H1 plate reader by collecting full light spectrum absorbance from 300 nm to 700 nm at 2 nm intervals. The absorbance at 730 nm was used to normalize the pigment content.

### Pigment quantification

A previously published protocol was adopted with some modifications^53^. 1.5 mL of mid-log phase cyanobacterial cultures were collected by centrifugation at 15,000 rpm for 5 min. Cell pellet was resuspended in 1.5 mL 90 % (Vol/Vol) cold acetone by vortexing. Cells were transferred to 2mL Bead-Beating DuraTubes and subjected to 2-3 cycles of bead-beating for 45 seconds each, with cooling on ice between cylces. After cell lysis, samples were centrifuged again for 2 min at 15,000 rpm. Supernatant were aliquoted onto 96-well microplate for measurement of A_665_ and A_720_ on the plate reader. Chlorophyll *a* was calculated as follows:

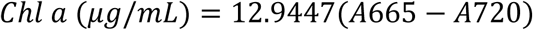

### Phylogenetic analyses of pD1

pD1 amino acid sequences of cyanobacteria, cyanophages, and red-lineage algae were downloaded from the UniProt Archive on 29 July 2024. First, the non-redundant database was searched using the term, “psbA”, and length criteria of 354-360 residues. This latter parameter was selected to minimize the incorporation of green algal or plant pD1 sequences, which typically have shorter tails and a total length of 353 residues. The initial search resulted in 2,177 unique sequences. Next, D1 capable of assembling the oxygen-evolving complex were selected by screening for essential residues D170, E189, E333, D342, and A344. This analysis resulted in 1,803 presumably active D1 sequences. These sequences were initially aligned using Clustal Omega. Low quality sequences with multiple unidentified residues or those with green lineage tails were manually culled resulting in 1,761 sequences. The Clustal Omega alignment was repeated, and the database analyzed using Geneious Prime version 2024.0.7. An unrooted phylogenetic tree was assembled using the Geneious tree builder package, which is a neighbor-joining method utilizing the Jukes-Cantor model for calculating genetic distances. For clarity, subtrees with phylogenetic distance ≤ 0.15 were collapsed.

## Acknowledgements

We thank Kuei-Yueh Ko for providing valuable insights in developing the statistical model used for screening sucrose producing mutants. We thank Daniel Ducat for gifting the plasmid construct used for constructing the sucrose-secreting cyanobacteria. We thank Duke Cancer Institute for gifting some supplies used for preliminary screening. We thank Julie Maupin-Furlow for carefully reading the manuscript. X.W. would like to acknowledge support from both the University of Florida and the Center for Bioinformatics and Functional Genomics at Miami University. Phylogenetic analyses were supported by U.S. Department of Energy, Office of Science, Office of Basic Energy Science, Division of Chemical Sciences, Geosciences, and Biosciences, Photosynthetic Systems through Grant DE-SC0020119 to D.J.V..

## Author contributions

X.W. conceived the work and drafted the manuscript. Z.J. designed and performed the third round high-throughput screening experiment, phenotype validation, genome sequencing analysis, and drafted the manuscript. K.N.I, M.W., and M.B contributed to several preliminary screening and validation experiments. X.Z. constructed the mutator strain and generated mutants for the first round of screening. D.W., A.F. and D.J.V conducted the evolutionary sequence analyses and drafted the related contents; M.O. provided valuable statistics insights for the validation of candidates from high-throughput screening; A.K. provided technical support for the high-throughput screening experiment. X.L. and all other authors reviewed and edited the manuscript.

## Data availability

All data are presented in the main text and supplemental materials.

## References

1 Zhu, X.-G., Long, S. P. & Ort, D. R. What is the maximum efficiency with which photosynthesis can convert solar energy into biomass? Curr. Opin. Biotechnol. 19, 153–159 (2008). 10.1016/j.copbio.2008.02.004

2 Boyer, J. S. Plant Productivity and Environment. Science 218, 443–448 (1982). doi:10.1126/science.218.4571.443

3 Wijffels, R. H. & Barbosa, M. J. An Outlook on Microalgal Biofuels. Science 329, 796–799 (2010). doi:10.1126/science.1189003

4 Long, S. P., Zhu, X.-G., Naidu, S. L. & Ort, D. R. Can improvement in photosynthesis increase crop yields? Plant Cell Environ. 29, 315–330 (2006). 10.1111/j.1365-3040.2005.01493.x

5 Ort, D. R. et al. Redesigning photosynthesis to sustainably meet global food and bioenergy demand. Proc. Natl. Acad. Sci. U. S. A. 112, 8529–8536 (2015). doi:10.1073/pnas.1424031112

6 Mertens, J. et al. Carbon capture and utilization: More than hiding CO_2_ for some time. Joule 7, 442–449 (2023). 10.1016/j.joule.2023.01.005

7 Falkowski, P. G. & Chen, Y.-B. in Light-Harvesting Antennas in Photosynthesis (eds Beverley R. Green & William W. Parson) 423-447 (Springer Netherlands, 2003).

8 De Souza, A. P. et al. Soybean photosynthesis and crop yield are improved by accelerating recovery from photoprotection. Science 377, 851–854 (2022). doi:10.1126/science.adc9831

9 Zhang, H., Zhao, Y. & Zhu, J.-K. Thriving under Stress: How Plants Balance Growth and the Stress Response. Dev. Cell 55, 529–543 (2020). 10.1016/j.devcel.2020.10.012

10 Gururani, M. A., Venkatesh, J. & Tran, L. S. P. Regulation of photosynthesis during abiotic stress-induced photoinhibition. Mol. Plant 8, 1304–1320 (2015).

11 Lawlor, D. W. & Tezara, W. Causes of decreased photosynthetic rate and metabolic capacity in water-deficient leaf cells: a critical evaluation of mechanisms and integration of processes. Ann. Bot. 103, 561–579 (2009).

12 Zhang, H., Li, Y. & Zhu, J.-K. Developing naturally stress-resistant crops for a sustainable agriculture. Nat. Plants 4, 989–996 (2018).

13 Clauw, P. et al. Leaf growth response to mild drought: natural variation in Arabidopsis sheds light on trait architecture. Plant Cell 28, 2417–2434 (2016).

14 Bechtold, U., Ferguson, J. N. & Mullineaux, P. M. To defend or to grow: lessons from Arabidopsis C24. J. Exp. Bot. 69, 2809–2821 (2018).

15 Shoval, O. et al. Evolutionary Trade-Offs, Pareto Optimality, and the Geometry of Phenotype Space. Science 336, 1157–1160 (2012). doi:10.1126/science.1217405

16 López-Calcagno, P. E. et al. Stimulating photosynthetic processes increases productivity and water-use efficiency in the field. Nat. Plants 6, 1054–1063 (2020). 10.1038/s41477-020-0740-1

17 Miyagawa, Y., Tamoi, M. & Shigeoka, S. Overexpression of a cyanobacterial fructose-1,6-/sedoheptulose-1, 7-bisphosphatase in tobacco enhances photosynthesis and growth. Nat. Biotechnol. 19, 965-969 (2001). 10.1038/nbt1001-965

18 Dann, M. et al. Enhancing photosynthesis at high light levels by adaptive laboratory evolution. Nat. Plants 7, 681–695 (2021). 10.1038/s41477-021-00904-2

19 Sun, H. et al. Engineered hypermutation adapts cyanobacterial photosynthesis to combined high light and high temperature stress. Nat. Commun. 14, 1238 (2023). 10.1038/s41467-023-36964-5

20 Negrão, S., Schmöckel, S. M. & Tester, M. Evaluating physiological responses of plants to salinity stress. Ann. Bot. 119, 1–11 (2016). 10.1093/aob/mcw191

21 Allakhverdiev, S. I. et al. Salt Stress Inhibits the Repair of Photodamaged Photosystem II by Suppressing the Transcription and Translation of *psbA* Genes in *Synechocystis*. Plant Physiol. 130, 1443–1453 (2002). 10.1104/pp.011114

22 Zörb, C., Geilfus, C.-M. & Dietz, K.-J. Salinity and crop yield. Plant Biol. 21, 31–38 (2019). 10.1111/plb.12884

23 Mori, T., Binder, B. & Johnson, C. H. Circadian gating of cell division in cyanobacteria growing with average doubling times of less than 24 hours. Proc. Natl. Acad. Sci. U. S. A. 93, 10183–10188 (1996). doi:10.1073/pnas.93.19.10183

24 Emlyn-Jones, D., Price, G. D. & Andrews, T. J. Nitrogen-regulated hypermutator strain of *Synechococcus* sp. for use in in vivo artificial evolution. Appl. Environ. Microbiol. 69, 6427–6433 (2003).

25 Hagemann, M. in Adv. Bot. Res. Vol. 65 (eds Franck Chauvat & Corinne Cassier-Chauvat) 27-55 (Academic Press, 2013).

26 Stitt, M., Herzog, B. & Heldt, H. W. Control of Photosynthetic Sucrose Synthesis by Fructose 2,6-Bisphosphate: I. Coordination of CO_2_ Fixation and Sucrose Synthesis. Plant Physiol. 75, 548-553 (1984). 10.1104/pp.75.3.548

27 Santos-Merino, M. et al. Improved photosynthetic capacity and photosystem I oxidation via heterologous metabolism engineering in cyanobacteria. Proc. Natl. Acad. Sci. U. S. A. 118 (2021). 10.1073/pnas.2021523118

28 Griese, M., Lange, C. & Soppa, J. Ploidy in cyanobacteria. FEMS Microbiol. Lett. 323, 124–131 (2011). 10.1111/j.1574-6968.2011.02368.x

29 Shen, J.-R. The Structure of Photosystem II and the Mechanism of Water Oxidation in Photosynthesis. Annu. Rev. Plant Biol. 66, 23–48 (2015). 10.1146/annurev-arplant-050312-120129

30 Ohad, I., Kyle, D. J. & Arntzen, C. J. Membrane protein damage and repair: removal and replacement of inactivated 32-kilodalton polypeptides in chloroplast membranes. J. Cell Biol. 99, 481–485 (1984). 10.1083/jcb.99.2.481

31 Satoh, K. & Yamamoto, Y. The carboxyl-terminal processing of precursor D1 protein of the photosystem II reaction center. Photosynth. Res. 94, 203–215 (2007). 10.1007/s11120-007-9191-z

32 Komenda, J. et al. Cleavage after residue Ala352 in the C-terminal extension is an early step in the maturation of the D1 subunit of Photosystem II in *Synechocystis* PCC 6803. Biochim. Biophys. Acta 1767, 829–837 (2007). 10.1016/j.bbabio.2007.01.005

33 Inagaki, N., Yamamoto, Y. & Satoh, K. A sequential two-step proteolytic process in the carboxyl-terminal truncation of precursor D1 protein in *Synechocystis* sp. PCC68031. FEBS Lett. 509, 197-201 (2001). 10.1016/S0014-5793(01)03180-5

34 Kuviková, S., Tichý, M. & Komenda, J. A role of the C-terminal extension of the photosystem II D1 protein in sensitivity of the cyanobacterium *Synechocystis* PCC 6803 to photoinhibition. Photochem. Photobiol. Sci. 4, 1044–1048 (2005). 10.1039/b506059a

35 Abramson, J. et al. Accurate structure prediction of biomolecular interactions with AlphaFold 3. Nature 630, 493–500 (2024). 10.1038/s41586-024-07487-w

36 Bekker, A. et al. Dating the rise of atmospheric oxygen. Nature 427, 117–120 (2004).

37 Croce, R. et al. Perspectives on improving photosynthesis to increase crop yield. Plant Cell (2024). 10.1093/plcell/koae132

38 Faralli, M. & Lawson, T. Natural genetic variation in photosynthesis: an untapped resource to increase crop yield potential? Plant J. 101, 518–528 (2020). 10.1111/tpj.14568

39 Clapero, V., Arrivault, S. & Stitt, M. Natural variation in metabolism of the Calvin-Benson cycle. Semin. Cell Dev. Biol. 155, 23–36 (2024). 10.1016/j.semcdb.2023.02.015

40 Ungerer, J., Wendt, K. E., Hendry, J. I., Maranas, C. D. & Pakrasi, H. B. Comparative genomics reveals the molecular determinants of rapid growth of the cyanobacterium *Synechococcus elongatus* UTEX 2973. Proc. Natl. Acad. Sci. U. S. A. (2018). 10.1073/pnas.1814912115

41 Golden, S. S., Brusslan, J. & Haselkorn, R. Expression of a family of *psbA* genes encoding a photosystem II polypeptide in the cyanobacterium *Anacystis nidulans* R2. EMBO J. 5, 2789–2798 (1986). 10.1002/j.1460-2075.1986.tb04569.x

42 Ivleva, N. B., Shestakov, S. V. & Pakrasi, H. B. The Carboxyl-Terminal Extension of the Precursor D1 Protein of Photosystem II Is Required for Optimal Photosynthetic Performance of the Cyanobacterium *Synechocystis* sp. PCC 68031. *Plant Physiol.* **124**, 1403-1412 (2000). 10.1104/pp.124.3.1403

43 Schrader, S. & Johanningmeier, U. The carboxy-terminal extension of the D1-precursor protein is dispensable for a functional photosystem II complex in Chlamydomonas reinhardtii. Plant Mol. Biol. 19, 251–256 (1992). 10.1007/BF00027346

44 Lers, A., Heifetz, P. B., Boynton, J. E., Gillham, N. W. & Osmond, C. B. The carboxyl-terminal extension of the D1 protein of photosystem II is not required for optimal photosynthetic performance under CO_2_- and light-saturated growth conditions. J. Biol. Chem. 267, 17494–17497 (1992). 10.1016/S0021-9258(19)37068-1

45 Sheridan, K. J., Duncan, E. J., Eaton-Rye, J. J. & Summerfield, T. C. The diversity and distribution of D1 proteins in cyanobacteria. Photosynth. Res. 145, 111–128 (2020). 10.1007/s11120-020-00762-7

46 Zurawski, G., Bohnert, H. J., Whitfeld, P. R. & Bottomley, W. Nucleotide sequence of the gene for the *M_r_* 32,000 thylakoid membrane protein from *Spinacia oleracea* and *Nicotiana debneyi* predicts a totally conserved primary translation product of *M_r_* 38,950. Proc. Natl. Acad. Sci. U. S. A. 79, 7699–7703 (1982). doi:10.1073/pnas.79.24.7699

47 Chen, J.-H. et al. Nuclear-encoded synthesis of the D1 subunit of photosystem II increases photosynthetic efficiency and crop yield. Nat. Plants 6, 570–580 (2020). 10.1038/s41477-020-0629-z

48 Andrews, S. FastQC: a quality control tool for high throughput sequence data. (2010). <http://www.bioinformatics.babraham.ac.uk/projects/fastqc/>.

49 Bolger, A. M., Lohse, M. & Usadel, B. Trimmomatic: a flexible trimmer for Illumina sequence data. Bioinformatics 30, 2114–2120 (2014). 10.1093/bioinformatics/btu170

50 Langmead, B. & Salzberg, S. L. Fast gapped-read alignment with Bowtie 2. Nat. Methods 9, 357–359 (2012).

51 Danecek, P. et al. Twelve years of SAMtools and BCFtools. GigaScience 10 (2021). 10.1093/gigascience/giab008

52 Seemann, T. Snippy: fast bacterial variant calling from NGS reads. (2018). <https://github.com/tseemann/snippy>.

53 Zavřel, T., Sinetova, M. A. & Červený, J. Measurement of Chlorophyll a and Carotenoids Concentration in Cyanobacteria. Bio. Protoc. 5, e1467 (2015). 10.21769/BioProtoc.1467

