## Supplementary information for "Enhancing Photosynthesis under Salt Stress via Directed Evolution in Cyanobacteria"

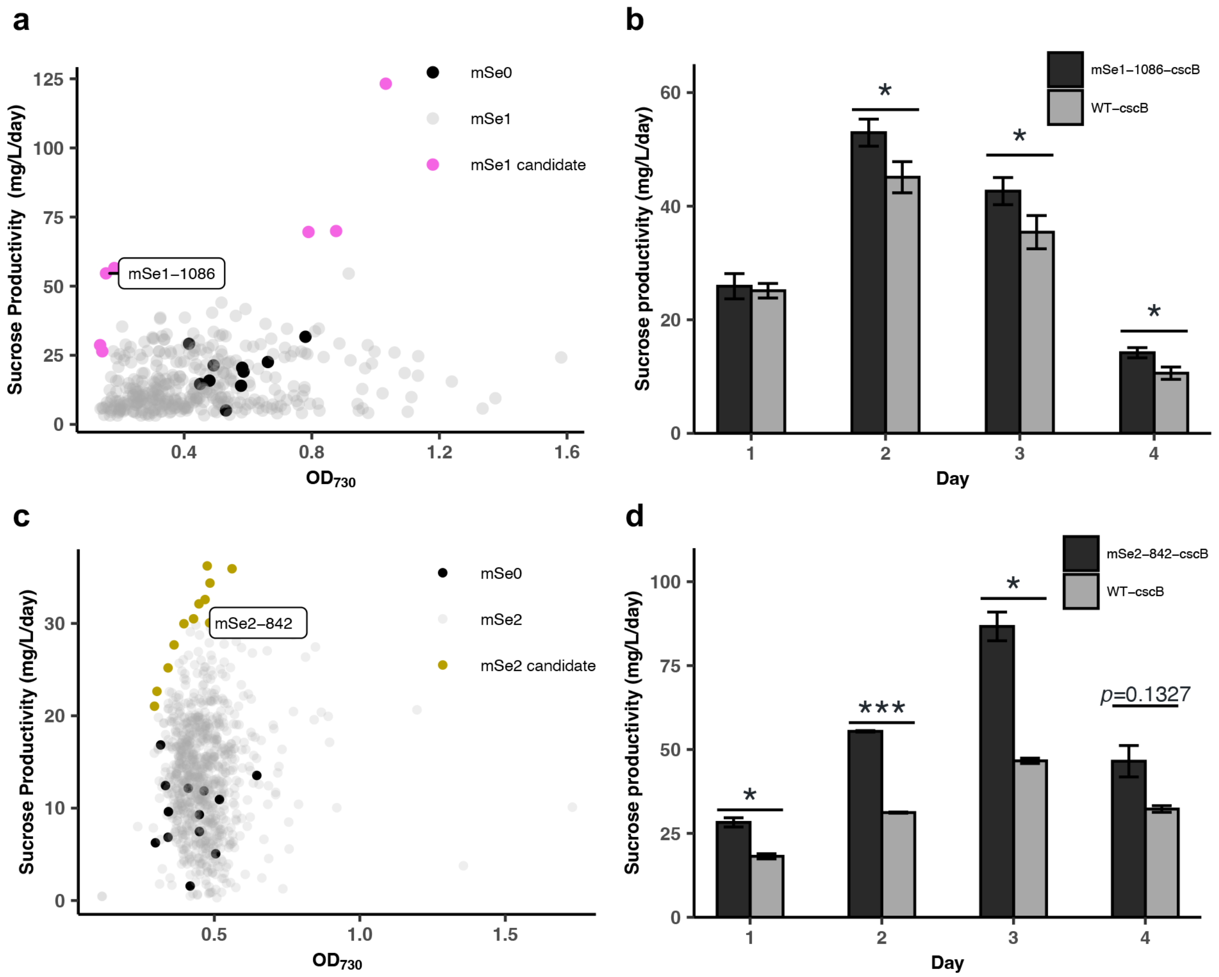


**Supplementary Figure 1 Sucrose production of mSe1 and mSe2 strains under salt stress conditions. a,** Sucrose productivities of mSe1 mutants after 18 hours of induction with 125 mM NaCl. Sucrose analysis excluded strains with no productivity. The strain mSe1-1086 was confirmed as the top sucrose producer after validation of the top candidate strains. **b,** Secreted sucrose in mSe1-1086_cscB and WT_cscB strains after induction with 150 mM NaCl with 1 mM IPTG. Error bars represent SD from three biological replicates. **c,** Sucrose productivities of mSe2 mutants after 30 hours of induction with 150 mM NaCl. The strain mSe2-842 was confirmed as the top sucrose producer after validation. **d,** Secreted sucrose in mSe2-842-cscB and WT_cscB strains after induction with 150 mM NaCl with 1 mM IPTG. Error bars show SD for two biological and technical replicates. Statistical significance is based on a one-tailed Student's t-test, with *$p < 0.05$, ***$p < 0.001$.

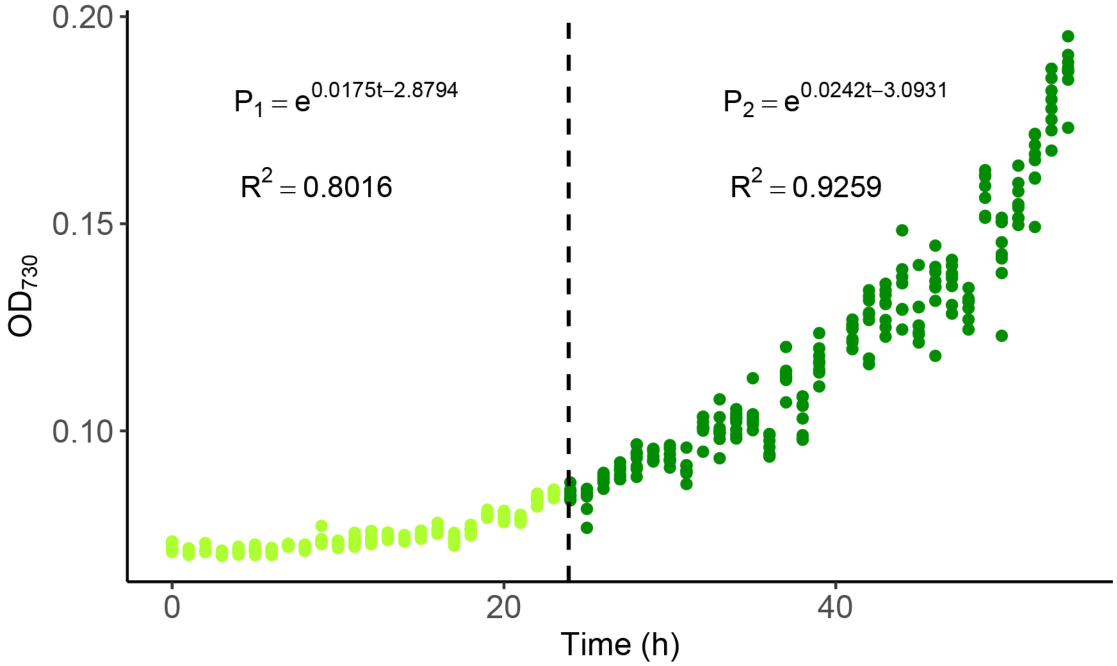


**Supplementary Figure 2 Growth of mSe0 strains in high-throughput screening system.** Growth was monitored in 96-well microplates, with data collected hourly. Salt induction occurred at 24 hours (indicated by the dashed line), with continuous data collection until 54 hours. Growth rates before and after salt induction were calculated using a linear model after log transformation of $OD_{730nm}$ with their respective adjusted $R^2$ shown in the figure.

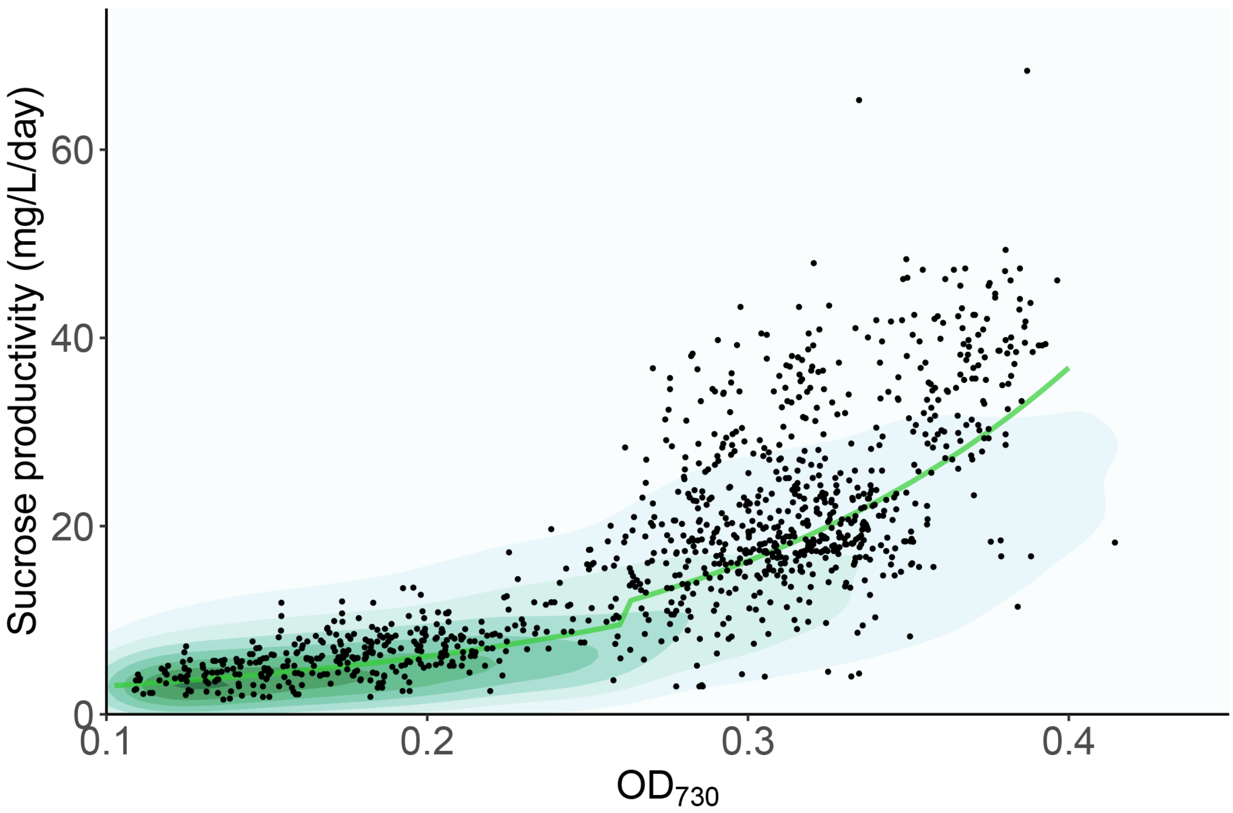


**Supplementary Figure 3 A segmented regression model to estimate the relationship between sucrose productivity and biomass in mSe0.** This scatter plot displays the sucrose productivity (mg/L/day) against biomass ($OD_{730nm,54h}$) for approximately 1,000 mSe0 samples. Data points are plotted alongside a fitted linear regression line (green), demonstrating the correlation between biomass and the logarithmic transformation of sucrose productivity. The shaded area represents the probability density, indicating the distribution and concentration of data points.

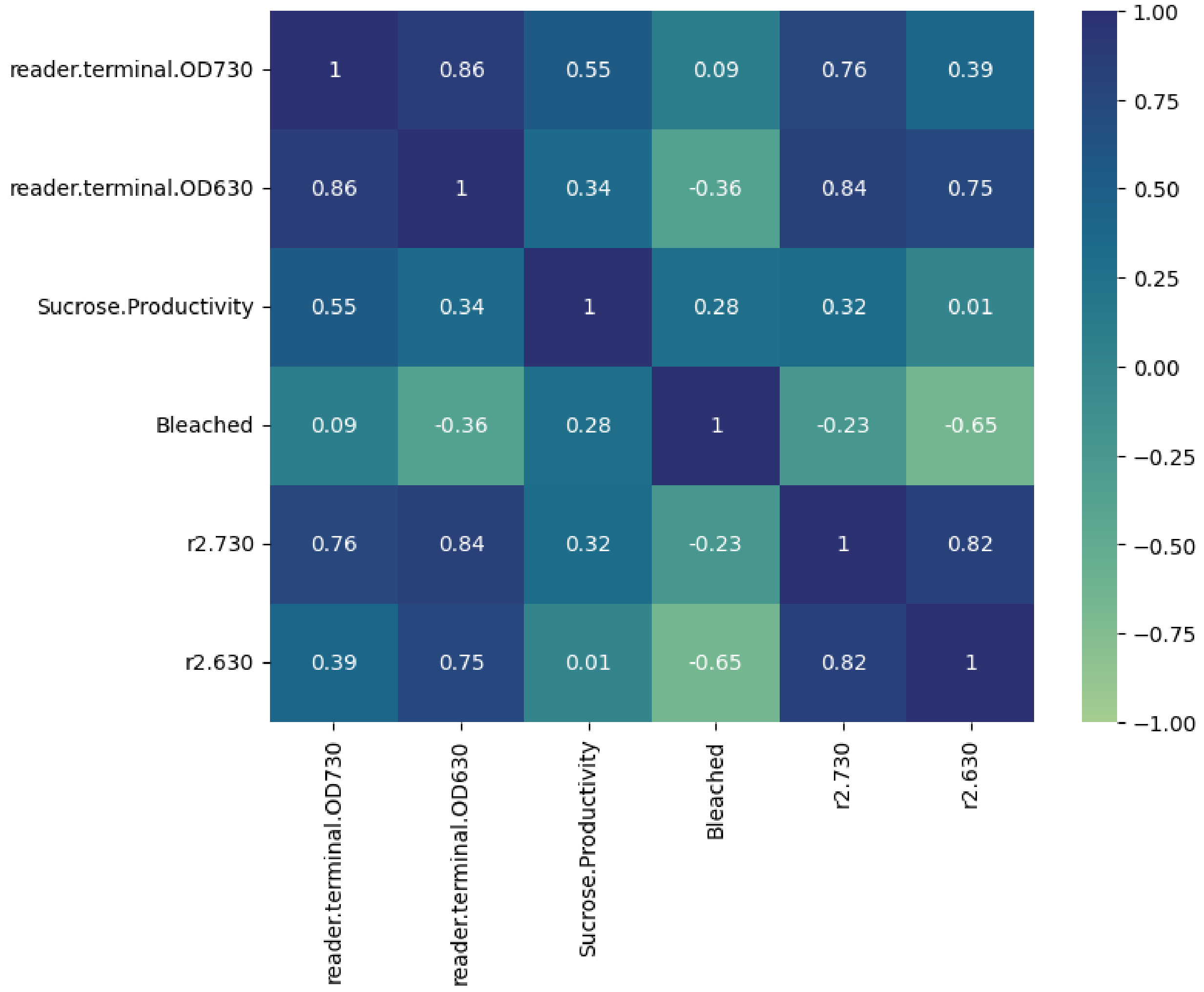


**Supplementary Figure 4 Correlation analysis.** $OD_{630nm,54h}$, $OD_{730nm,54h}$, sucrose productivity, growth rate post-salt induction ($OD_{730nm,54h}/OD_{730nm,24h}$), and phycobilin pigment change rate ($OD_{630nm,54h}/OD_{630nm,24h}$) were included as different variables and tested against each other and against the chlorosis status. reader.termninal.OD730: $OD_{730nm,54h}$, reader.termninal.OD630: $OD_{630nm,54h}$, Sucrose.Productivity: sucrose productivity, Bleached: chlorosis status, r2.730: growth rate represented by the division of $OD_{730nm,54h}$ and $OD_{730nm,24h}$, r2.630: phycobilin pigment change rate represented by the division of $OD_{630nm,54h}$ and $OD_{630nm,24h}$.

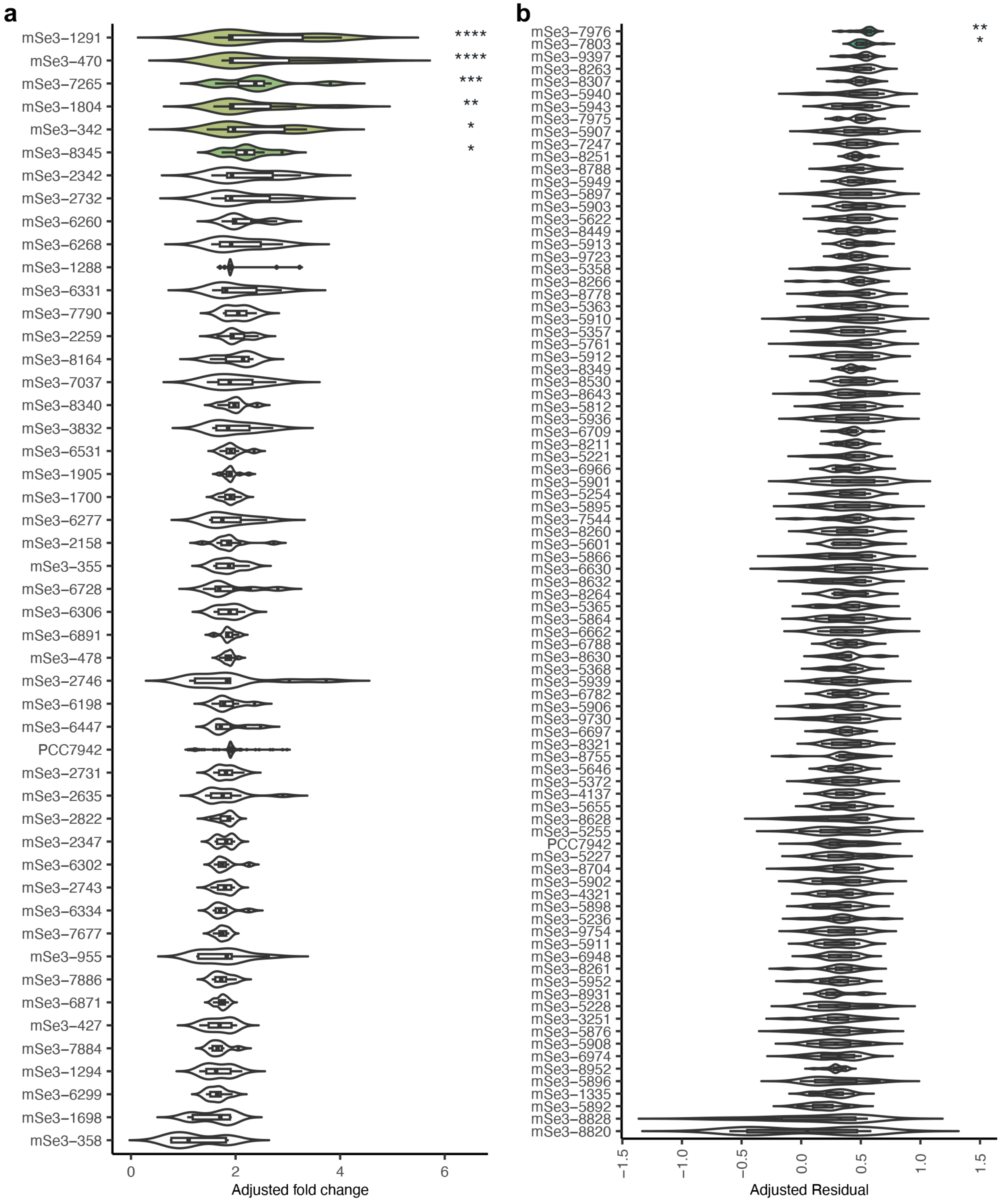


**Supplementary Figure 5 Validation for biomass and sucrose productivity of top candidate strains from high-throughput screening.** Following the determination of proper sample size through power analysis, replicates of each candidate from three different classes (**a**: BAM and FGM; **b**: SPM) were inoculated into 96-well microplates under the same condition as preliminary screening. For biomass validation, 12 replicates (4 replicates × 3 batches) reached the power of

89.7% to detect an effect size difference of 0.1 OD. For sucrose validation, 16 replicates (8 replicates × 2 batches) reached the power of 98% to detect an effect size difference of 25 residual sucrose productivity. The plots were based on adjusted biomass fold change or sucrose residuals after normalization. To remove batch effect, block ANOVA was performed with the block term representing different batches. **a,** mSe3-1291, mSe3-470, mSe3-7265, mSe3-1804, mSe3-342, mSe3-8345 showed statistical differences using block ANOVA. **b,** mSe3-7976 and mSe3-7803 showed statistical differences using block ANOVA. (*$p < 0.05$, ** $p < 0.01$, *** $p < 0.001$, **** $p < 0.0001$).

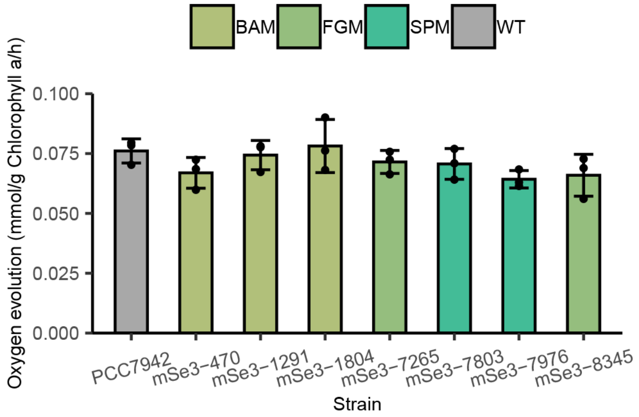


**Supplementary Figure 6 Oxygen evolution measurement for elite BAM, FGM, and SPM strains in multi-cultivators.** The maximum $O_2$ evolution rates of the mutants are similar to the wild type *S. elongatus* PCC 7942, suggesting normal photosynthetic efficiency of these mutants.

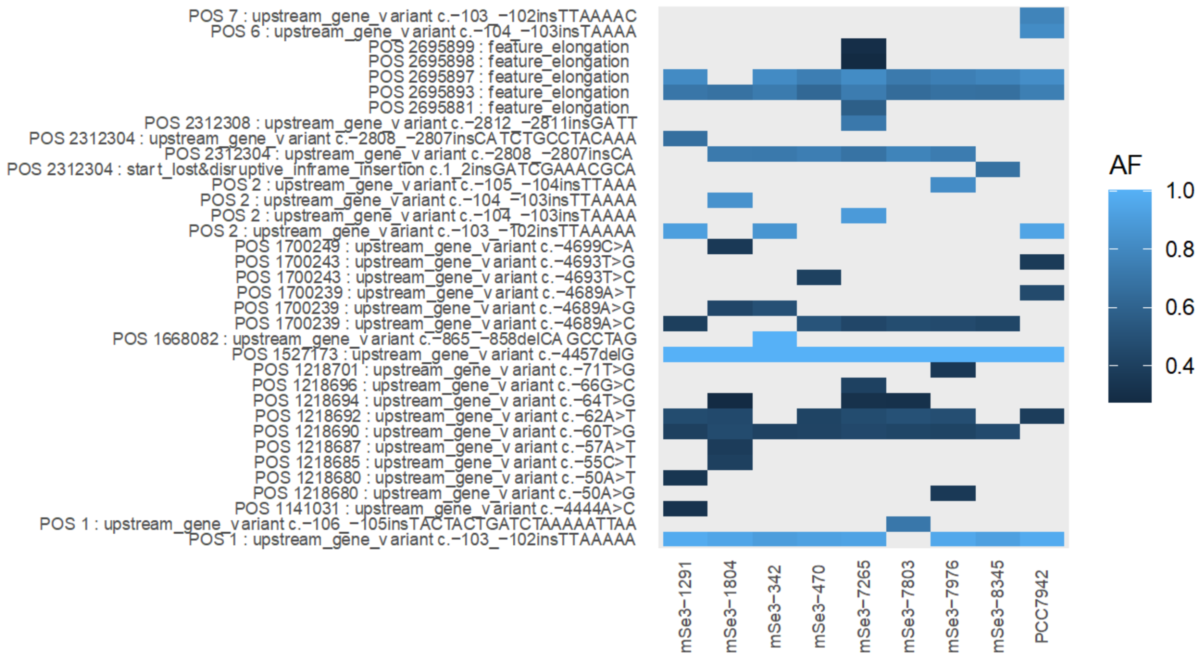


**Supplementary Figure 7 Mutations in non-coding DNA (ncDNA) with allele frequencies above 0.25.** Mutations are indicated with the position followed by respective mutations. SNPs are represented by $N_1$>$N_2$, with each $N_i$ being one of the four nucleotides. Insertion mutations are annotated as “ins” and followed by inserted nucleotides, and deletions are annotated as “del” followed by deleted nucleotide(s). The negative number led by “c.” represents the location upstream of the nearest CDS. Notably, a deletion in ncDNA results in a filamentous phenotype in mSe3-342.

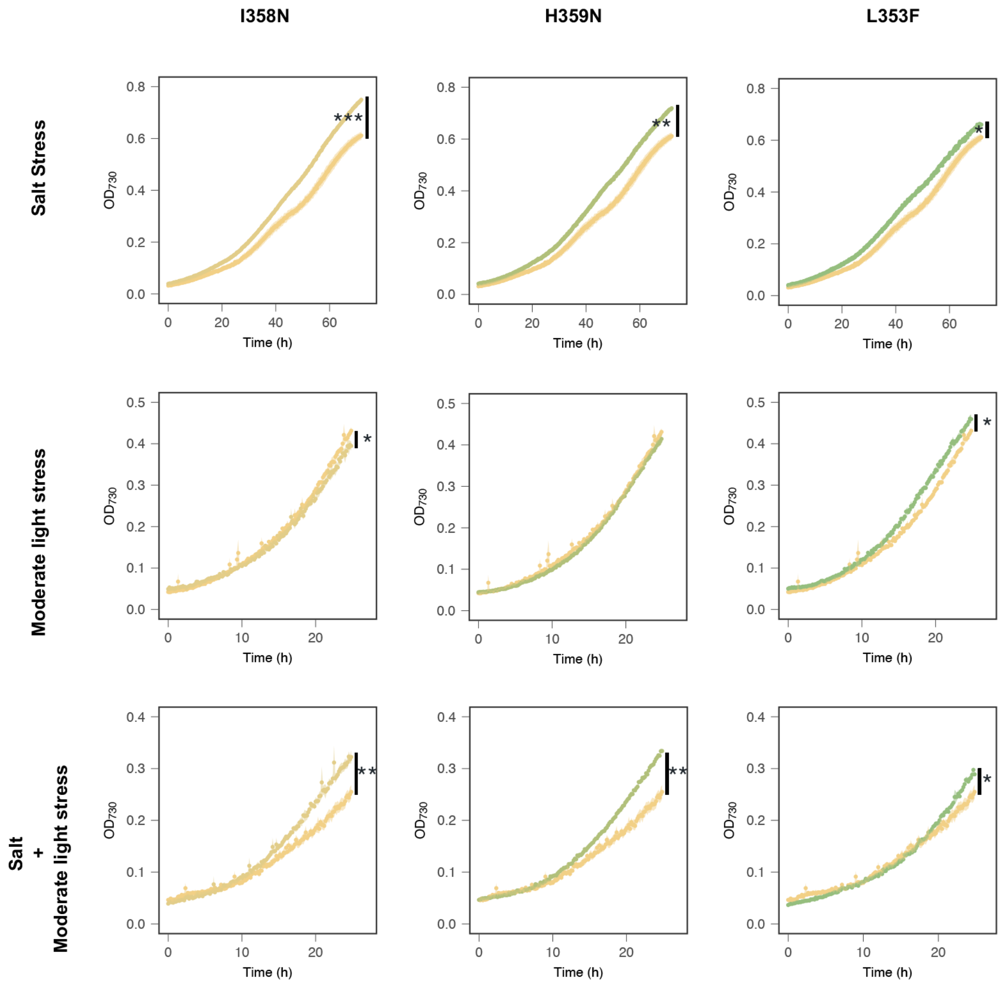


**Supplementary Figure 8 Growth dynamics of transformants with mutations in *psbA1* under stress conditions.** First row (salt stress, 150mM NaCl): Mutants L353F, I358N, and H359N exhibited increased biomass than *psbA1*$^{WT}$. Second row (moderate light stress, 300 μE·m$^{-2}$·s$^{-1}$): Mutants L353F and I358N showed slightly higher biomass compared to *psbA1*$^{WT}$. Third row (combined stress): All mutants showed higher biomass yield than the *psbA1*$^{WT}$ control. Statistical significance was assessed using an unpaired Student's t-test (*$p < 0.05$, ** $p < 0.01$, *** $p < 0.001$).

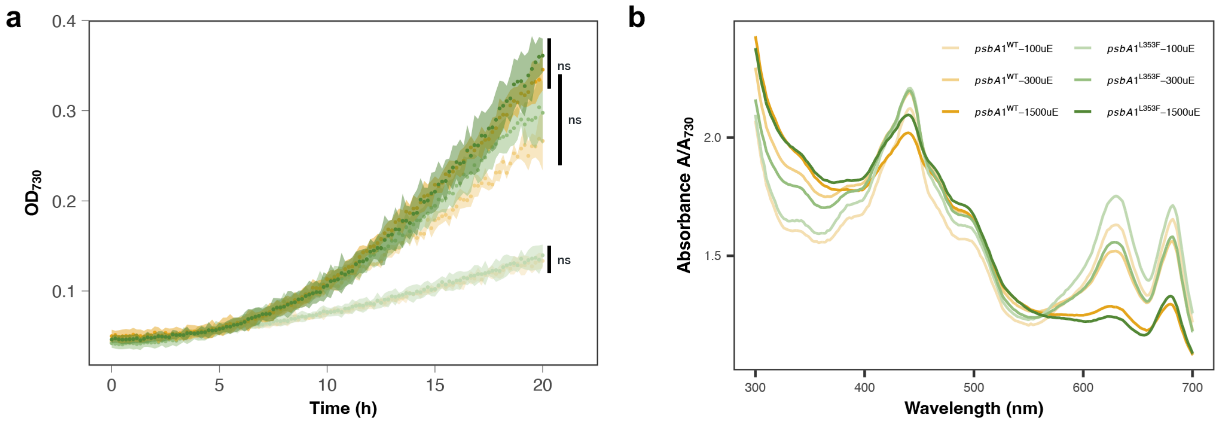


**Supplementary Figure 9 Growth and full light spectrum scanning of *psbA1*$^{WT}$ and L353F mutant. a,** Growth dynamics. No significant growth difference between the *psbA1*$^{WT}$ and L353F mutant was observed under light intensities of 100 (top curves), 300 (middle curves), and 1,500 (bottom curves) $\mu E \cdot m^{-2} \cdot s^{-1}$. **b,** Light absorption spectrum. At 1,500 $\mu E \cdot m^{-2} \cdot s^{-1}$, the L353F mutant showed lower phycobilin ($OD_{630nm}$) but higher chlorophyll contents ($OD_{440nm}$), with the opposite shown under lower light intensities.

**Supplementary Table 1 Primers for RT-qPCR analysis.**

| Primer | Sequence 5’-3’ |
|---|---|
| ***mutS*** | F- CATTCAACTGGCTCAAA |
| | R- TTCGTTCATCCACCATGA |
| ***rnpB*** | F- ACCAGACTTGCTGGGTAACG |
| | R- TTACCGAGCCAACACCTCTC |
| ***ppc*** | F- AATCTTGATTCGGCCATCTG |
| | R- AGAAAGCCCTAGGGACGGTA |
| ***secA*** | F- GCTATGGGCAAAAAGATCCA |
| | R- AACCACCTCAGCTTGTGGAC |

**Supplementary Table 2 Mutations on *psbA1* CDS.**

| Strain | Mutation | Location on genome |
|---|---|---|
| mSe3-470 | His359Asn | 414947 |
| mSe3-7803 | His359Asn | 414947 |
| mSe3-7976 | Thr354Pro (AF < 0.25) | 414932 |
| | Ile358Asn | 414945 |
| | His359Asn | 414947 |
| mSe3-8345 | Leu353Phe | 414931 |
| | Thr354Ala (AF < 0.25) | 414932 |
| | Ile358Asn | 414945 |

**Supplementary Table 3 Primers for mutation recovery confirmation.**

| Strain | Sequence 5'-3' |
|---|---|
| All | F: atgtctagATGACCAGCATTCTTCGCGAGCAACGCCGCGA |
| *psbA1*$^{WT}$ | R: atcGGATCCTTAACCGTGAATTGAAGGCGCAGTCAAAGCGACCGGGGTCGCT |
| I358N | R: atcGGATCCTTAACCGTGAtTTGAAGGCGCAGTCAAAGCGACCGGGGTCGCT |
| H359N | R: atcGGATCCTTAACCGTtAATTGAAGGCGCAGTCAAAGCGACCGGGGTCGCT |
| L353F | R: atcGGATCCTTAACCGTGAATTGAAGGCGCAGTaAAAGCGACCGGGGTCGCT |
| T354P+I358N+H359N | R: atcGGATCCTTAACCGTtAtTTGAAGGCGCAGgCAAAGCGACCGGGGTCGCT |
| L353F+T354A+I358N | R: atcGGATCCTTAACCGTGAtTTGAAGGCGCAGcaAAAGCGACCGGGGTCGCT |

### Supplementary Table 4 Distribution of amino acids in pD1 tail sequences.

| | 345 | 346 | 347 | 348 | 349 | 350 | 351 | 352 | 353 | 354 | 355 | 356 | 357 | 358 | 359 | 360 |
|---|---|---|---|---|---|---|---|---|---|---|---|---|---|---|---|---|
| A | 49.3% | 14.0% | 3.1% | 24.6% | 37.0% | 0.5% | 0.7% | 95.0% | 0.0% | 3.9% | 94.9% | 0.4% | 63.3% | 0.1% | 3.7% | 5.4% |
| C | 0.7% | 0.0% | 0.0% | 0.3% | 0.8% | 0.0% | 0.0% | 0.0% | 0.0% | 0.0% | 0.1% | 0.0% | 0.0% | 0.0% | 0.0% | 0.0% |
| D | 0.1% | 1.0% | 18.5% | 0.0% | 0.0% | 0.0% | 0.0% | 0.0% | 0.0% | 0.2% | 0.2% | 0.1% | 0.4% | 0.0% | 0.6% | 0.0% |
| E | 0.1% | 0.6% | 70.0% | 2.4% | 1.5% | 0.1% | 0.0% | 0.0% | 0.0% | 0.1% | 0.0% | 0.0% | 1.3% | 0.0% | 0.8% | 0.0% |
| F | 0.3% | 0.1% | 0.0% | 1.6% | 0.2% | 0.0% | 0.0% | 0.0% | 2.9% | 0.0% | 0.0% | 0.0% | 0.0% | 0.0% | 0.1% | 0.0% |
| G | 0.8% | 63.0% | 0.2% | 0.1% | 0.0% | 0.0% | 0.0% | 0.3% | 0.0% | 1.2% | 0.2% | 0.2% | 0.1% | 0.2% | 0.3% | 86.8% |
| H | 0.0% | 0.1% | 0.0% | 0.0% | 0.1% | 0.0% | 0.0% | 0.0% | 0.0% | 0.1% | 0.1% | 0.0% | 0.1% | 0.0% | 7.2% | 0.0% |
| I | 0.1% | 0.3% | 0.1% | 2.9% | 2.6% | 0.1% | 4.9% | 0.8% | 6.0% | 5.0% | 0.0% | 0.1% | 0.4% | 82.0% | 0.5% | 0.1% |
| K | 0.0% | 0.0% | 0.5% | 0.0% | 0.7% | 0.0% | 0.0% | 0.0% | 0.2% | 1.0% | 0.3% | 0.0% | 0.3% | 0.0% | 0.3% | 0.0% |
| L | 0.1% | 1.0% | 0.1% | 0.6% | 22.5% | 0.5% | 5.7% | 0.1% | 72.7% | 0.3% | 0.0% | 0.2% | 0.5% | 1.1% | 0.1% | 0.0% |
| M | 0.2% | 0.1% | 0.0% | 0.6% | 1.1% | 0.3% | 0.1% | 0.0% | 12.2% | 0.3% | 0.0% | 0.0% | 0.0% | 0.3% | 0.1% | 0.0% |
| N | 0.2% | 4.4% | 2.6% | 0.1% | 0.1% | 0.1% | 0.0% | 0.5% | 0.1% | 3.4% | 0.3% | 0.0% | 0.3% | 0.0% | 70.8% | 0.0% |
| P | 0.0% | 0.0% | 0.5% | 2.4% | 0.0% | 94.8% | 0.0% | 0.2% | 0.0% | 0.0% | 0.1% | 96.8% | 0.2% | 0.0% | 0.0% | 0.0% |
| Q | 0.0% | 4.3% | 1.0% | 3.8% | 4.9% | 0.3% | 0.0% | 0.0% | 0.4% | 6.9% | 0.1% | 0.1% | 2.3% | 0.0% | 1.8% | 0.0% |
| R | 0.1% | 0.1% | 0.1% | 0.0% | 0.3% | 0.4% | 0.0% | 0.0% | 0.4% | 0.7% | 0.1% | 0.0% | 0.2% | 0.0% | 0.1% | 0.0% |
| S | 44.6% | 5.2% | 1.1% | 25.3% | 1.0% | 0.8% | 0.1% | 1.3% | 0.2% | 16.6% | 1.3% | 0.5% | 20.4% | 0.1% | 1.4% | 5.6% |
| T | 1.7% | 2.6% | 1.2% | 3.2% | 21.7% | 1.2% | 0.2% | 0.1% | 0.3% | 43.8% | 0.9% | 0.1% | 2.6% | 0.1% | 0.6% | 0.0% |
| V | 1.7% | 3.2% | 1.0% | 32.1% | 4.9% | 0.3% | 87.1% | 0.5% | 3.4% | 16.2% | 0.3% | 0.3% | 6.0% | 14.5% | 1.1% | 0.2% |
| W | 0.0% | 0.0% | 0.0% | 0.0% | 0.0% | 0.0% | 0.0% | 0.0% | 0.2% | 0.0% | 0.0% | 0.0% | 0.0% | 0.0% | 0.0% | 0.0% |
| Y | 0.0% | 0.0% | 0.0% | 0.0% | 0.5% | 0.0% | 0.0% | 0.0% | 0.1% | 0.0% | 0.0% | 0.1% | 0.1% | 0.0% | 0.3% | 0.0% |
| gap | 0.0% | 0.0% | 0.0% | 0.0% | 0.3% | 0.7% | 1.2% | 1.2% | 0.9% | 0.4% | 1.2% | 1.3% | 1.6% | 1.6% | 10.4% | 2.0% |
